# The NOX2-ROS-NLRP3 inflammasome axis in traumatic brain injury

**DOI:** 10.1101/2025.05.16.654460

**Authors:** Janeen Laabei, Gloria Vegliante, Nathan Ryzewski Strogulski, Carly Douglas, Sahil Threja, Andrew Pearson, Aurore Nkiliza, Marie Hanscom, Isabella Dias Filogonio Emediato, Fiona Crawford, Joseph Ojo, David J. Loane

**Author notes:** Address correspondence to: David Loane Ph.D., School of Biochemistry and Immunology, Trinity College Dublin, Room 5.08C, 152-160 Pearse Street, Dublin, D02R590, Ireland. **Data availability declaration:** Data are available upon request and can be accessed through the corresponding author. **Ethics, consent to participate, and consent to publish declarations**: Not applicable.

## Abstract

**Background:** Phagocyte NADPH oxidase 2 (NOX2) is an enzyme complex responsible for reactive oxygen species (ROS) production. Chronic NOX2 activity sustains oxidative stress/damage and drives neuroinflammation following traumatic brain injury (TBI). NOX2 acts as a priming signal for NLRP3 inflammasome activation, which also plays a role in secondary injury after TBI. GSK2795039 is a small molecule brain penetrable drug that inhibits NOX2 in a NADPH competitive manner. Here, we investigated whether pharmacological inhibition of NOX2 using GSK2795039 can reduce secondary neuroinflammation after TBI, specifically via inhibition of downstream NLRP3 inflammasome activation, in both resident microglia and infiltrating myeloid cells in the injured brain.

**Methods:** Immortalised microglial (IMG) cells or primary microglia were pre-treated with GSK2795039 (NOX2 inhibitor) or MCC950 (NLRP3 inhibitor) and stimulated with lipopolysaccharide and nigericin to induce NOX2/ROS and NLRP3 inflammasome activation. The controlled cortical impact model, pharmacokinetic analyses, multi-dimensional flow cytometry, histology and neurobehavioral assessments were used to translate *in vitro* findings to an experimental TBI model in adult male C57BL6/J mice.

**Results:** The small molecule NOX2 inhibitor, GSK2795039, attenuated microglial NOX2 activity, thereby reducing ROS, nitrite and cytokine levels, as well as NLRP3 inflammasome components in pro-inflammatory microglia. TBI recruited NOX2/ROS/IL-1β+ neutrophils and inflammatory monocytes into injured brain parenchyma with peak monocytic NOX2/ROS/IL-1β production at 3 days post-injury (DPI), coincident with peak NOX2/ROS/IL-1β expression in microglia. Systemic administration of GSK2795039 (100mg/kg; I.P.) starting 2 hours post-injury attenuated NOX2/IL-1β+ microglial and infiltrating myeloid cell activation at 3 DPI. In addition, NOX2 inhibition reduced numbers of IL-1R+ T cells in the brain of TBI mice, indicating that myeloid-T cell crosstalk was altered by GSK2795039 treatment. Innate and adaptive neuroimmune changes were associated with improvements in motor function post-TBI. In the chronic phase through 28 DPI, pharmacological inhibition of NOX2 by GSK2795039 treatment resulted in modest improvements in neurobehavioral deficits and TBI neuropathology.

**Conclusions:** These preclinical studies identify the NOX2-ROS-NLRP3 inflammasome axis along with myeloid-T cell crosstalk as effective targets for TBI neuroinflammation. Our translational studies indicate that NOX2 may be a promising therapeutic target for mitigating neuroinflammation in microglia, and peripheral immune cells, following experimental TBI in mice.

## Introduction

Phagocyte NADPH oxidase 2 (NOX2) is a multi-subunit enzyme system responsible for the generation of intracellular and extracellular reactive oxygen species (ROS) in immune cells. NOX2 activation stimulates reactive microgliosis in age-related neurodegenerative diseases, including Alzheimer’s disease (AD) and Parkinson’s disease (PD) [1], and it contributes to oxidative stress/damage in chronic traumatic brain injury (TBI) [2–5]. NOX2 is robustly increased in CD68+ reactive microglia within the lesion boundary [2–4], and it is also highly expressed in infiltrating F4/80+ inflammatory monocytes acutely following TBI [4], indicating peripheral and central (i.e. brain resident) mechanisms of action. Genetic ablation of NOX2 is neuroprotective in TBI and reduces oxidative stress/damage, decreases microglial-mediated neurotoxicity, while also promoting hippocampal neurogenesis and cognitive function recovery after TBI [4–7]. Further, NOX2 deletion upregulates anti-inflammatory mechanisms in microglia/macrophages, which are due, in part, to increased IL-10 and STAT3 signalling in the injured cortex and hippocampus [8]. Upregulated post-traumatic microglial/macrophage NOX2 expression is coincident with Nod-like receptor protein 3 (NLRP3) inflammasome activation and increased pro-inflammatory IL-1β production [9, 10].

Mechanistically, NOX2/ROS triggers NLRP3 inflammasome formation and activation in response to exogenous stimuli (PAMPs) as well as endogenously produced or secreted molecules from damaged cells (DAMPs) [11]. Inhibition of NOX2-derived ROS prevents ATP-induced caspase-1 activation and IL-1β production in alveolar macrophages [12], while knockdown of the p22^phox^ subunit of NOX2 significantly suppresses IL-1β release in THP1 cells in response to asbestos and MSU challenge [13]. Thioredoxin binding protein (TXNIP), one of the α-arrestin protein superfamily, is involved in this ROS-dependent inflammatory response [14]. In the resting state, TXNIP is kept inactive by binding with its endogenous inhibitor thioredoxin (TRX). Both TXNIP and TRX regulate and maintain the balance of intracellular oxidation and anti-oxidation systems. When TXNIP separates from TRX, it activates endoplasmic reticulum stress-mediated NLRP3 inflammasome complex formation, triggers mitochondrial stress-induced apoptosis, and stimulates inflammatory cell death (pyro ptosis) [14]. Thus, NOX2-dependent ROS is a priming signal for NLRP3 inflammasome activation, which results in caspase-1 dependent cleavage of IL-1β and IL-18. Notably, Ma et al. reported increased expression of TXNIP and activation of NLRP3 inflammasome in the injured cortex acutely after experimental TBI in mice, and that NOX2 knockout markedly attenuated NLRP3 inflammasome expression [15].

Clinically, NOX2 and NLRP3 inflammasome are activated in neurodegenerative diseases, such as AD [16, 17]. Further, TXNIP and IL-1β are overexpressed in the AD brain and associated with tau pathogenesis [18]. *In vitro* molecular studies that exposed microglia to pTau containing neuronal media, exosomes, and paired helical filaments, results in IL-1β activation in a NLRP3-ASC-caspase-1 dependent manner [19]. Moreover, intracerebral injections of homogenates containing fibrillar amyloid-β and tau-seeding competent aggregated tau activates the NLRP3-ASC inflammasome and induces tau propagation in an NLRP3 dependent manner [20, 21]. Notably, genetic or pharmacological manipulation of components within the NLRP3 inflammasome, reduce tau seeding, hyperphosphorylation and aggregation, and improves cognitive function in tau transgenic mice [20, 21]. In mouse models, TBI induces a self-propagating tau pathology that progressively spreads in the brain [22], while in chronic human TBI, dystrophic microglia have been shown to spatially co-exist with tau lesions in end-stage autopsy tissue [23, 24].

Our long-term goal is to identify molecular mechanisms causing posttraumatic neuroinflammation that contribute to progressive brain degeneration and associated neurological impairments (e.g. dementia). In this study we hypothesised that NOX2 is an upstream regulator of NLRP3 inflammasome activation in microglia, such that pharmacological inhibition of the NOX2/ROS/NLRP3 inflammasome axis using a small molecule NOX2 inhibitor, GSK2795039, will mitigate the deleterious effects of neuroinflammation on TBI outcomes. GSK2795039 is a novel brain penetrant and specific NOX2 inhibitor shown to reduce NOX2-dependent ROS *in vitro* and *in vivo* in a NADPH competitive manner [25]. We first investigated the inhibitory actions of GSK2795039 using *in vitro* models in microglia induced by lipopolysaccharide and nigericin that stimulate NOX2/ROS signalling and NLRP3 inflammasome activation, pro-inflammatory cytokine production (e.g. IL-1β, IL-18), and pyroptotic cell death. We compared the effects of NOX2 inhibition on the NOX2/ROS/NLRP3 inflammasome axis against MCC950, a potent NLRP3 inflammasome inhibitor that has been shown to suppress microglial mediated neuroinflammation and associated neurodegeneration in experimental TBI models in mice [26, 27]. We then translated our studies to the controlled cortical impact model in mice to determine whether pharmacological inhibition of NOX2 using GSK2795039 could disrupt the NOX2/ROS/NLRP3 inflammasome axis to reduce secondary neuroinflammation after TBI and improve long-term outcomes.

## Materials and Methods

### Materials

GSK2795039 (CAS No.: 1415925-18-6; CAT# HY-18950) was purchased from Medchem Express, adenosine 5’- triphosphate disodium salt hydrate (ATP; CAS No.: 34369-07-8; CAT# A26209), lipopolysaccharide (LPS, E. Coli 055:B5; CAS No.: 93572-42-0; CAT# L6529) and MCC950 (CAS No.: 256373-96-3; CAT# 5381200001) from Sigma-Aldrich, and nigericin (CAS No.: 28643-80-3; CAT# tlrl-nig) from InvivoGen. Primary antibodies were purchased as indicated: anti-NLRP3 (AdipoGen Life Sciences; Cat# AG-20B-0014-C100, RRID:AB_2885199), anti-Casapse-1 (AdipoGen; Cat# AG-20B-0042-C100, RRID:AB_2755041), anti-ASC (AdipoGen; Cat# AG-25B-0006-C100, RRID:AB_2885200), and anti-β-actin (Sigma-Aldrich; Cat# A1978, RRID:AB_476692). HRP-conjugated secondary antibodies were purchased from Jackson Immuno Research Laboratories. For flow cytometry, fluorophore conjugated primary antibodies were purchased as indicated: Brilliant violet-785 conjugated anti-mouse CD45 (BioLegend; Cat# 103149, RRID:AB_2564590), APC-eFluror-780 conjugated anti-mouse CD11b (Thermo Fisher Scientific; Cat# 47-0112-82, RRID:AB_1603193), Brilliant Violet-421 conjugated anti-mouse Ly6G (BioLegend; Cat# 127627, RRID:AB_10897944), Brilliant Violet–605 conjugated anti-mouse Ly6C (BD Biosciences; Cat# 563011, RRID:AB_2737949), Pe-Dazzle-594 conjugated anti-mouse CD3 (BioLegend; Cat# 100245, RRID:AB_2565882), PE-Cynanine5 conjugated anti-mouse CD4 (Thermo Fisher Scientific; Cat# 15-0041-82, RRID:AB_468695), Alexa-Fluor-700 conjugated anti-mouse CD8 (Thermo Fisher Scientific; Cat# 56-0081-82, RRID:AB_494005), CD121a (IL-1RI) Rat anti-mouse, PE, clone:35F5 (BD Biosciences; Cat # 557489, RRID: AB_396727), PerCP-eFluor 710 conjugated anti-mouse IL-1β (Thermo Fisher Scientific; Cat# 46-7114-82, RRID: AB_2573835), Alexa Fluor 647 conjugated anti-mouse NOX2/gp91^phox^ (Bioss; Cat# bs-3889R-A647-BSS), Dihydrorhodamine 123 (DHR123; Invitrogen; Cat # D23806),Live/Dead Fixable AQUA (Invitrogen; Cat # L34957).

### Cell culture

*A) Immortalised microglia (IMG) cell line:* IMG cells (Kerafast; Cat# EF4001, RRID:CVCL_HC49; [28]) were cultured in complete Dulbecco’s Modified Eagle’s Medium (cDMEM) containing 4.5g/L glutamine (Sigma-Aldrich) supplemented with 10% heat-inactivated fetal bovine serum (FBS; Sigma-Aldrich), 1% (v/v) penicillin/streptomycin (Sigma-Aldrich) at 37°C with 5% CO2. The IMG cells were cultured in T75 flasks for 3-4 days until fully confluent and cells between passage 5-20 were used for experimentation. Once confluent, cells were trypsinised using Trypsin-EDTA (Sigma-Aldrich; 5min, 37°C), centrifuged (1350 RPM; 5min, room temperature) and the pelleted cells were resuspended in cDMEM. Cell solution was counted using Trypan Blue (Sigma-Aldrich). IMG cells were seeded at a density of 7.5 x 10^4^ cells/well (96-well) and 1x10^6^ cells/well (12-well).

*B) Primary microglia:* Primary microglia were cultured from cerebral cortices of postnatal day 1 Wistar Rat pups.

The brains were isolated, and the cerebral cortices were placed in a petri-dish containing DMEM/F12 (Gibco) and freed from meninges. The cortices were washed once in DMEM/F12 and trypsinised (0.25% Trypsin/EDTA, 1X; Invitrogen) for 10min at 37°C, followed by triturating (repeated pipetting approximately 20 times) in growth medium (DMEM/F12 with 10% FBS and 1% (v/v) penicillin-streptomycin) to dissociate tissue. Cells were collected by centrifugation (1,000 RPM; 10min, room temperature), resuspended in DMEM/F12 and the mixed glial suspension were plated in Poly-D-Lysine (Sigma-Aldrich) coated T75 culture flasks at 37°C with 5% CO2. The medium was replaced following 24h, and after 7-10 days in culture the microglia were detached from the mixed glial culture using an orbital shaker (Incu-Shaker mini, Benchmark Scientific; 100 RPM; 1h, 37°C), centrifuged (1000 RPM; 10min, room temperature) and cells were counted using Trypan blue. Microglia were plated in 96-well plates at a density of 1x10^5^ cells/well in DMEM/F12 medium (Gibco) supplemented with 10% FBS and 1% (v/v) penicillin-streptomycin.

*C) In vitro treatments:* IMGs and primary microglia were pre-treated with GSK2795039 (1-20µM; 1h, 37°C) or MCC950 (0.005-0.5 µM; 1h, 37°C), stimulated with lipopolysaccharide (LPS; 100ng/ml; 8h), followed by an additional stimulation with nigericin (10µM; 30min).

### Plate-based assays

*A) Intracellular ROS:* Intracellular ROS was measured using 5-(and-6)-chloromethyl-2’,7’- dichlorodihydrofluoresceindiacetate (CM-H2DCFDA; Invitrogen; Cat #C6827). Following stimulation, cells were incubated with CM-H2DCFDA for 45min at 37°C in 5% CO2, as per manufacturer’s instruction. Fluorescence was measured immediately at excitation and emission wavelengths of 485 and 528nm, respectively, using Spectra Max Gemini Microplate Reader. (Molecular Devices, USA). Data are presented as a percentage of controls.

*B) Nitrite release: A* Griess reagent assay (Invitrogen; Cat# G7921) was used to measure nitrite release into the supernatant, according to manufacturer’s instruction. The absorbance was read at 548 nm using Versa MAX tuneable Microplate Reader with SoftMax Pro software (Molecular Devices, USA). Unknown samples were extrapolated using a standard curve of sodium nitrite (NaNO2; 2.5-20µM).

*C) Cell viability:* Immediately following the ROS assay, cell viability was assessed using a microculture (3-(4,5- dimethylthiazol-2-yl)-2,5-diphenyltetrazolium bromide (MTT; Sigma-Aldrich; Cat# M2128)-based colorimetric assay, according to manufacturer’s instructions, using Versa MAX tuneable Microplate Reader with SoftMax Pro software (Molecular Devices, USA). Data are presented as a percentage of controls.

*D) Lactate dehydrogenase (LDH):* Lactate dehydrogenase (LDH) was measured in the supernatant as an indicator of pyroptosis using the CytoTox 96® Non-Radioactive Cytotoxicity Assay (Promega; Cat# G1780), according to manufacturer’s instructions. Data are presented as a percentage of controls.

*E) Cytokines release:* Cytokine levels in cell culture supernatant were quantified by enzyme-linked immunosorbent assay using TNFα and IL-1β specific assays (R&D Systems; Cat# DY410 and DY401 respectively), according to the manufacturer’s instructions. Cytokine levels were expressed as pg of cytokines/ml.

***NOX activity:*** Microglial NADPH oxidase activity was quantified using lucigenin-enhanced chemiluminescence [29]. Briefly, IMG cells in 12-well plates were harvested in 400μL ice-cold PBS, lifted using a sterile cell scraper, and homogenised using a 1mL micropipette. Total superoxide production was assessed in 100μL of cell suspension, transferred to white 96 well microplates containing NADPH (100μM; TCI Europe; Cat# T0521) and lucigenin (100μM; TCI Europe; Cat# 2315971). Wells containing SOD (15IU; Sigma-Aldrich; Cat# S5395) for each sample were used to subtract the SOD-uninhibitable portion from the total superoxide production. Additionally, a well containing the NOX inhibitor, apocynin (2mM; Sigma-Aldrich; Cat# W508454), was added as a negative control. Relative luminescence was measured at 37°C for 1h using a Luminoskan™ Ascent plate reader (ThermoFisher). NOX-dependent superoxide production was calculated as the fold of control of the integrated area under the curve of SOD-inhibitable lucigenin chemiluminescence.

***Western immunoblot:*** IMGs were cultured in 12-well plates, washed with ice-cold PBS and protein was isolated using radioimmunoprecipitation assay buffer (RIPA) containing protease and phosphatase (phosphatase inhibitor cocktail 2 & 3; Sigma-Aldrich) for 20min on ice. Cells were sonicated on ice and centrifuged (15,000 RPM; 15min, 4°C) to pellet the insoluble protein. The cell lysates (supernatant) were collected, and protein concentration was determined using Pierce^TM^ BCA Protein Assay kit (Thermo Scientific; Cat# 23227). Samples were normalised to 1µg/µl using RIPA and loaded onto precast 8-15% SDS-polyacrylamide gels (Bio-Rad). To blot for supernatants, the cultured medium from IMG treated cells were collected and StrataClean resin (Agilent; Cat# 400714) was added (1:100 dilution), vortexed and rotated overnight at 4°C. The following day the resin was collected by centrifugation (5,000x g, 5min, 4°C), supernatant discarded, with the captured proteins bound to the resin beads. The resin was resuspended in SDS-PAGE sample buffer (60µl/tube containing β-mercaptoethanol (1:10) and boiled at 99°C for 5min). The resin was pelleted by centrifugation, and the sample (20µl) was loaded onto precast 8-15% SDS-polyacrylamide gels (Bio-Rad). Cell lysates of equal protein or supernatants of equal volume were electrophoresed through 8-15% SDS-polyacrylamide gels. The gels were transferred to 0.2μM pore-sized nitrocellulose membranes using Trans-Blot® Turbo™ Transfer Starter System (Bio-Rad) followed by blocking in 5% milk in TBS-Tween-20 (TBST). After blocking, the membranes were incubated with primary antibodies (diluted in 5% TBST), including anti-NLRP3 (1:1000), anti-Caspase-1 (1:1000), anti-ASC (1:1000), anti-IL-1β (1:1000), anti-β-actin (1:50,000) overnight at 4°C. The membranes were washed in TBST (3 x 5min) and incubated in HRP-conjugated secondary antibodies in TBST for 2h at room temperature. Finally, the membranes were washed (3 x 5min) and detection was carried out using enhanced chemiluminescence (WesternBright^TM^ ECL-spray; Advansta). Proteins were normalised to β-actin and quantified using Image Lab software v6.0.1 build 34, where data are expressed in arbitrary units.

***Controlled cortical impact:*** Mice were used in compliance with EU regulations as overseen by the Health Products Regulatory Authority (HPRA), Ireland. All animal experiments and surgical procedures were designed in accordance with the Animals in Research: Reporting *In vivo* Experiments (ARRIVE) guidelines and protocols were approved by the Trinity College Dublin Animal Research Ethics Committee and HPRA (Protocol Number: AE19136/P138). Adult male C57BL/6J mice (10–12-week-old; 24-27g) were bred by the Comparative Medicine Unit (CMU) in Trinity College Dublin. All mice were housed in groups of five per cage in specific pathogen-free conditions (12h light/dark cycle, 24±1°C and 55±5% humidity) with *ad libitum* access to food and water. Controlled cortical impact (CCI) was performed using the precise impactor device (RWD; Cat# 6809II) as described previously [30]. Following induction of anaesthesia with isoflurane (3% induction, 0.75% for maintenance using O2 flow rate), Bupivacaine (0.083%) was administered subcutaneously near the site of incision. The head of the mouse was mounted in a stereotaxic frame. The surgical site was cleaned with ethanol and chlorohexidine and a 10mm midline incision was made and the skin reflected. A 5mm craniotomy was made on the central aspect of the left parietal bone while maintaining surgical anaesthesia and sterile conditions. Moderate-level brain injury was induced using a 3mm bevelled impactor tip at 3m/sec velocity, 1.5mm deformation depth, and 0.18m/sec dwell time. Immediately following CCI, bupivacaine (0.083%) was administered at the injury site, the incision was closed with interrupted 5-0 silk sutures, and saline (1ml) was administered subcutaneously on the dorsal side of the mouse. Sham animals underwent the same procedure as injured mice except for craniotomy and cortical impact. The sutures were cleaned daily for three consecutive days and animals were monitored throughout the study using HPRA approved score sheets. For *in vivo* drug treatment studies, GSK2795039 (MedChem Express), was prepared in 10% DMSO and 90% Corn Oil on the day of the injections. Vehicle (10% DMSO, 90% Corn Oil) served as control. Randomly assigned mice were administered GSK2795039 intraperitoneally (i.p) after CCI (first injection at 2h post-injury), with repeated injections delivered by a blinded researcher.

**Drug biodistribution analyses by LC/MS:** A) Plasma preparation: Briefly, 50μL of plasma was mixed with 20μL of 100μM ND630 in 100% LC/MS grade methanol (used as internal standard) and 130μL of 100% LC/MS grade methanol. Samples were then vortexed, left on wet ice for 30min, and later centrifuged for 10min at 10,800RPM and 4°C. Supernatants were collected and filtered using the PVDF 0.2µM centrifuge filters, and flow through were used to make 200μl of a 1:100 dilution in 75% LC/MS grade methanol and transferred to chromatography autosampler vials for LC/MS analyses. B) Cortex, Hippocampus and Liver preparation: Briefly, all three types of tissue were homogenized with 750μL of 100% LC/MS grade methanol using a Dounce homogenizer (total volume ∼1mL). 2μL of 100μM ND630 (for liver/cortex) and 2μL of 10μM ND630 (for hippocampus) in 100% LC/MS grade methanol (used as internal standard) was added to the homogenate, vortexed, and centrifuged at 20,000x g for 10min. Supernatants were collected and filtered using the PVDF 0.2µM centrifuge filters at 20,000x g for 5min. Flow through were transferred into an autosampler vial and dried using a speed-vac. Dried samples of Cortex and Liver were reconstituted/homogenized with 2ml of 75% LC/MS grade methanol. While dried samples of the hippocampus were reconstituted/homogenized with 200μl of 75% LC/MS grade methanol. 200μl of reconstituted samples from all three types of tissue were transferred into a chromatography autosampler vial with inset for LC/MS analyses.

*LC/MS analyses:* An Ultimate 3000 LC system interfaced to an Orbitrap Exploris 480 mass spectrometer was used for LC–MS/MS analysis. Samples were separated using a Acquity UPLC BEH C18, 1x50mm, 1.7μm LC column (Waters), where solvent A contained 10mM of ammonium formate and 0.1% of formic acid and solvent B consisted of 100% acetonitrile. Separation was performed by isocratic elution for 8min with a flow rate at 50μL/min, where the mobile phase composition remained at 45% B throughout the run. All samples were kept at 4°C in the auto-sampler tray for the duration of the analysis. Data was acquired in positive mode using full scan with a mass range of m/z 400–600 and resolution of 60,000. For increased sensitivity, data were also acquired using positive mode with single ion monitoring for GSK2795039 and ND-630 (used as internal standard); m/z 451.191 ± 4 and 570.191 ± 4, respectively, at a resolution of 60,500. Peak areas of GSK2795039 and ND-630 were obtained from Tracefinder™ at a window of five parts per million accuracy. GSK2795039 concentrations were calculated based on relative spiked internal standard amount and interpolated from a standard curve ranging from 5nM to 150nM. Each sample was injected in triplicate, and all triplicate runs with a CV above 20% were excluded from further analysis.

**Neurobehavioral testing:** Mice were handled for 5min/day for a minimum of 5 days prior to starting the experiments, and prior to testing mice were acclimatized for 30-60min in the behaviour room. For all tests, even illumination (35 lux, confirmed by a luminometer (Extech LT300 Light Meter)) and constant noise (45dB generated by a white noise machine; confirmed by a decibel meter (Extech 407730 Digital Sound Level Meter)) was used in the behaviour room. Neurobehavioral tests were video-tracked using ANY-maze cameras and software (Stoelting Europe).

*Beam walk:* Fine motor coordination was assessed using a beam walk test as previously described [31]. Briefly, mice were placed at the end of a wooden beam (5mm wide, 1300mm long; suspended 30-40cm above foam padding), and the number of foot faults of the right hind limb was recorded over 50 steps, as the mouse moved towards an escape house at the other end of the elevated beam. Mice were trained on the beam walk for 3 consecutive days prior to CCI. Mice with more than 10-foot faults did not meet the basic performance criteria for the beam walk test and were excluded from the final analysis.

*Two-trial Y-maze:* To evaluate spatial working memory, the two-trial Y-maze was performed as previously described [32, 33]. In the first trial, a randomised selected arm was blocked with a guillotine and the mouse was placed in a randomly selected starting arm, facing the wall. The mouse was allowed to freely explore the maze for 5min before being placed in the holding cage for an inter-trial-interval of 1h before starting trial 2. In the second trial, the guillotine was removed from the blocked arm, the mouse was placed in the same starting arm as in trial 1 and allowed to freely explore all 3 arms for 5min. The time the mice spent in the novel arm was determined.

*Accelerating rotarod:* Gross motor function was assessed with rotarod as previously described [34]. Briefly, mice were trained 3 days prior to sham/CCI and were tested each day post-injury. Mice were placed on an accelerating rotarod (Ugo Basile) that increased in speed from 4-60 RPM with maximum speed reached at 180sec. The latency to fall from the rod, or hold onto and rotate with the rod twice, was recorded for three trials per day. A 10min rest period with access to food and water was allowed between each trial. The scores were averaged from the three trials to give a single score per mouse per day.

*Novel object recognition (NOR)*: For assessment of recognition memory, the NOR test was used as previously described [31]. Mice underwent two habituation days consisting of one 10min trial/day to acclimate to the testing arena (22.5 x 22.5cm, black plexiglass walls). Twenty-four hours following the second habituation, mice underwent object familiarisation in which mice were placed into the arena containing two identical objects, equidistance apart, and allowed to freely explore the objects for 5min. Twenty-four hours later, mice underwent novel object testing whereby one object was replaced with a novel object. Mice were allowed to freely explore the objects for 5min. The novel object was equally distributed across experimental groups, and the side the novel object was placed was randomised and balanced across groups to avoid object and side bias. The amount of time the mice spent exploring the objects were recorded manually using timers. Object interaction included sniffing and interacting with the object directly. Mice with a cumulative exploration time (both objects) of less than 6sec were excluded from the study. The discrimination index (DI) was calculated as follows: (time exploring novel – time exploring familiar) / (total time novel + familiar object). Mice with a DI of 0.2 are considered to have an intact cognitive function [31].

***Real-Time Quantitative Polymerase Chain Reaction (RT-qPCR):*** Quantitative gene expression analysis in the hippocampus of sham/CCI mice was performed using Taqman technology as previously described [9]. Briefly, total RNA was extracted from hippocampal tissue punches using RNeasy^®^ Mini kit (Qiagen) and cDNA was synthesized using Verso cDNA Synthesis Kit (ThermoFisher Scientific), as per manufacturer’s protocol. Real-time PCR amplification was carried out using 7500 Fast Real-Time PCR system (Applied Biosystems) with SensiFAST Probe Lo-ROX (Bioline; BIO-84020). The following Taqman assays were used: *nlrp3*, Mm00840904_m1; *tnf*, Mm00443258_m1; *il1b*, Mm00434228_m1; *cd68*, Mm03047343_m1; *cybb*, Mm01287743_m1; *cyba*, Mm00514478_m1; *gapdh*, Mm99999915_g1. Relative mRNA expression was calculated using the comparative cycle threshold (2^−ΔΔ^Ct) method, and GAPDH served as endogenous control. Relative mRNA expression is presented as fold change over sham control.

***Flow cytometry:*** Mice were humanely euthanised by isoflurane overdose and perfused through the left ventricle with 30ml cold PBS before the removal of the brain. The ipsilateral cortex, excluding olfactory bulb and cerebellum, were sectioned and collected into 1ml cRPMI. DNase and Collagenase (1mg/ml) were added into the tubes and incubated at 37°C for 1h on a shaker (Incu-Shaker mini, Benchmark Scientific). The homogenate was passed through a 70µM cell strainer, topped up with cRMPI, centrifuged and resuspended in 40% Percoll before being layered over 70% percoll and centrifuged with the brake off (600x g; 20min, 4°C). The upper layer of myelin was discarded, and the isolated mononuclear cells were collected from the interface of the Percoll gradient. Cells were washed with cRPMI and plated on a 96-well V-plate and immediately stained for flow cytometry. Following incubation with Brefeldin A (Sigma Aldrich; Cat #B7651) and Monensin (BD Biosciences; Cat # 554724) for 3h at 37°C, CountBright^TM^ Absolute Counting Beads (Invitrogen; Cat# C36950) were added, cells were centrifuged and stained with Live/Dead (Invitrogen; Cat# L34957; 15min; dark, room temperature). Cells were washed with FACS buffer and resuspended in a master mix of mouse Fc Block (BD Pharmingen; Cat# 553141) and surface fluor-conjugated antibodies (1:200-400 dilutions): CD45-BV785 (Cat# 103149), CD11b-APCeF780 (Cat # 47-0112-82), Ly6C-BV605 (Cat # 563011), Ly6G-BV421 (Cat # 127627), CD3-PeDazzle^594^ (Cat # 100245), CD4-PeCy-5 (Cat #15-0041-82), CD8-AF700 (Cat # 56-0081-82) and IL-1R-PE (Cat #557489). Following a wash step, cells underwent intracellular staining whereby cells were fixed by adding Fixation/Permeabilization working solution and incubated for 30min at room temperature. Cells were washed twice with 1X Permeabilization buffer and centrifuged (1350 RPM; 5min, 4°C). The pelleted cells were re-suspended in NOX2/gp91^phox^-AF647 (1:100, Cat# bs-3889R-A647) and IL-1β-PercP710 (1:200, Cat# 46-7114-82) in 1X Permeabilization buffer, and incubated overnight at 4°C (in dark). Cells were washed twice in 1X Permeabilization buffer and finally washed in FACS buffer for analysis. To measure ROS, isolated mononuclear cells from the ipsilateral cortex were processed into a single-cell suspension in 96-well V-plates and immediately stained with the ROS indicator DHR123 (Invitrogen; 1:500, 30min at 37°C). Cells were washed with PBS and stained with Live/Dead (Invitrogen; 50µl/tube diluted 1:600) in PBS and incubated (15min in dark at room temperature). Isolated mononuclear cells were then washed with FACS buffer and underwent surface staining using surface fluor-conjugated antibodies as described before. For *in vitro* flow cytometry studies, IMGs were cultured on 12-well plates and treated with drugs / LPS + nigericin prior to processing into a single-cell suspension for intracellular staining with NOX2/gp91^phox^-AF647, as described before. All flow cytometric data was acquired using Cytek Aurora Spectral flow cytometry with SpectroFlo^®^ software. Compensation was calculated using single stained cells. Cell-specific fluorescence minus one (FMO) controls were used for gating strategies and flow cytometric analysis was performed using FloJo software (TreeStar Inc.). Mean Fluorescence Intensity (MFI) of cells was measured on live, single cells, and cell populations were identified using the gating strategy shown in **supplemental figure 1**. The absolute number of cells were calculated based on the frequency of the total parent population.

***Lesion volume:*** Mice were perfused with saline (60ml), followed by 4% PFA (60ml) using a pump (Watson-Marlow 120S) to ensure a constant flow rate. The brain was isolated and placed in 4% PFA for a 3h post-fix. Next, brains were transferred into 30% sucrose for cryopreservation (up to 2 days), frozen using isopentane and liquid nitrogen, and stored in foil in -70°C until sectioning. Brains were sectioned in 40µm using Cryostat (Leica CM3050 S) and coronal slices were serially collected at from bregma +1.2mm to -3.6mm at 200µm apart, directly on glass slides (Superfrost^TM^ Plus Adhesion Microscope Slides; Epredia; J1800AMNZ) or in 24-well plates containing 0.01M PBS. This 5-series cutting scheme (two sections on two glass slides and the next three sections in individual 24-wells) allowed for a wide range of brain regions across the anterior-posterior axis. A total of 5 sections/well were stored in freezing solution (250ml PBS; 75g Sucrose; 75ml ethylene glycol) in -80°C until further analysis. Coronal brain sections were stained with Cresyl violet for lesion volume quantification. Briefly, slices were de-hydrated following incubation in 70% - 90%- 100% ethanol prior to incubation with Xylene to de-lipidise the tissue. Next the tissue was incubated in 100% - 90%-70% ethanol and finally H2O to fully hydrate the tissue prior to staining with Cresyl violet. The slices were dehydrated with acetic acid (3%) to de-stain and clean up parenchyma before mounting with DPX mounting medium. Images were acquired by Leica DM4 B microscope and LAS X software. Lesion volume quantification was analysed by Fiji software. The total volume (mm^3^) of the lesion on the ipsilateral side was calculated as the tissue loss compared to the contralateral side [35, 36].

**Statistical analysis:** Blinding within the study was performed as follows: 1) individual who administered GSK2795039 was blinded to injury group, 2) behavioural and stereological analyses were performed by individuals blinded to injury and treatment groups. Statistical analysis was performed using Prism v9.1.0 software (GraphPad software, San Diego, CA, USA; RRID: SCR_002798). Normality testing was performed using the Shapiro-Wilks test, and where data passed normality parametric statistical analysis was performed. Data was analysed by unpaired student’s *t* test, One-way ANOVA with post hoc Dunnett’s multiple comparisons test, or Two-Way ANOVA, with or without repeated measures with uncorrected Fisher’s LSD test. All quantitative data are represented as mean ± standard error of the mean (SEM). For *in vitro* studies, data represent examples of three independent experiments. p values < 0.05 were considered statistically significant.

## Results

### Selective inhibition of NOX2 using a small molecule inhibitor, GSK2795039, reduces NOX2 activity, ROS, nitrite, and TNF⍺ in pro-inflammatory microglia

To closely mimic what occurs in TBI, we established an *in vitro* model of microglial activation using LPS/nigericin (L/N) stimulation to induce both NOX2 and NLRP3 inflammasome activation in immortalised microglia cells (IMG; [28]). Nigericin induces potassium (K^+^) efflux resulting in pore formation of the plasma membrane to induce NLRP3 inflammasome activation [37, 38]. L/N potently induces pyroptotic dependent IL-1β release. We sought to determine whether GSK2795039, a small molecule NOX2 inhibitor [25], could attenuate NOX2-mediated pro-inflammatory activation of microglia *in vitro*. L/N increased the mean florescent intensity (MFI) of NOX2 protein in IMG microglia (**Figure 1A**; p<0.0001 vs control). L/N increased NOX-driven superoxide production (**Figure 1B**; p<0.01 vs control). GSK2795039 attenuated NOX2 MFI in a concentration-dependent manner (**Figure 1A**; p<0.0001 vs L/N), whereas MCC950 pre-treatment, an NLRP3 inhibitor, had no effect on NOX2 MFI (**Figure 1A**). GSK2795039 (10µM) and MCC950 reduced NOX activity in L/N simulated IMGs (**Figure 1B**; p<0.05; p<0.01 vs L/N). These findings demonstrate that GSK2795039 inhibits NOX2 protein expression and activity in a model of NLRP3 inflammasome activation in microglia. In addition, the potent NLRP3 inhibitor, MCC950, fails to reduce NOX2 expression.

**Figure 1:**
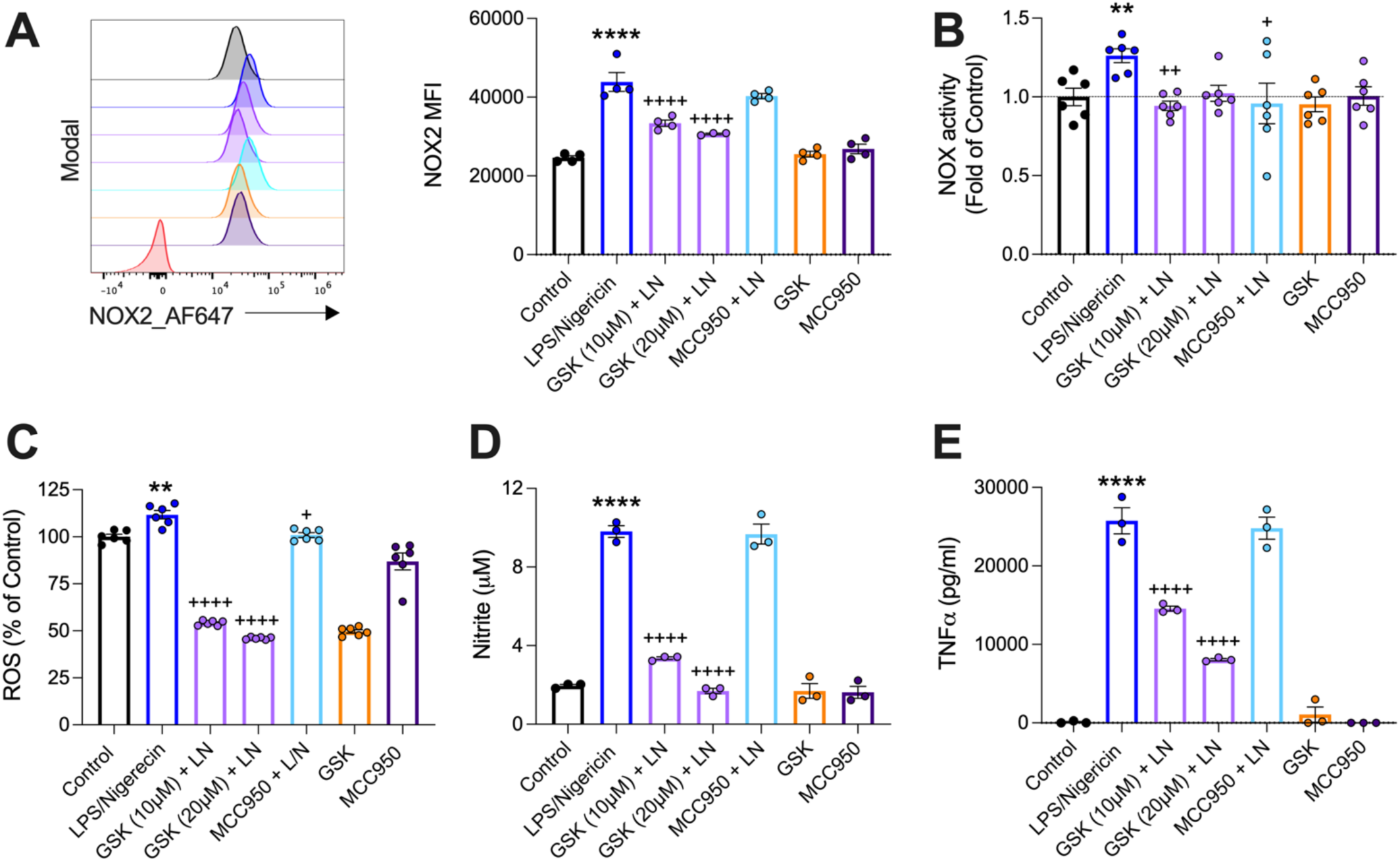
GSK2795039 attenuates NOX2 expression and activity, reactive oxygen species, nitrite, and TNF⍺ production in IMGs. IMGs pre-treated with GSK2795039 (10 or 20 µM) or MCC950 (0.5 µM) for 1h prior to stimulation with LPS/nigericin (L/N; 10 µM; 30 min) in a 12-well plate. GSK2795039 (10 µM) and MCC950 (0.01 µM) were used as drug controls. L/N increased NOX2 mean florescent intensity (MFI; A; ****p<0.0001), NOX2 activity and ROS (B,C; **p<0.01), nitrite and TNF⍺ (D,E; ****p<0.0001) compared to controls. GSK2795039 reduced NOX2 MFI (A; ^++++^p<0.0001), NOX activity (B; ^++^p<0.01), ROS, nitrite and TNF⍺ (C-E; ^++++^p<0.0001) versus L/N. MCC950 reduced superoxide activity and ROS (B, C; ^+^p<0.05) versus L/N but had no effect on NOX2 MFI (A), nitrite (D) or TNF⍺ (E). Data are mean ± SEM (n=3-6 per group) and are a representative of three independent experiments. **p<0.01; ****p<0.0001 versus control, ^+^p<0.05; ^++^p<0.01; ^++++^p<0.0001 versus L/N; One-way ANOVA with post hoc Dunnett’s multiple comparisons test versus L/N.

We next investigated what effect inhibition had on the pro-inflammatory mediators downstream of the NOX2/ROS signalling pathway. In microglia, ROS production induces NFκB translocation and NOS2 (iNOS) activation, followed by release of nitric oxide [39, 40]. Moreover, ROS and IL-1β results in an overproduction of TNF⍺ that can promote caspase-1 activity and drive neuroinflammation [41]. Therefore, we investigated the effect of GSK2795039 and MCC950 on ROS, nitrite and TNF⍺ levels pro-inflammatory microglia. L/N increased ROS, nitrite and TNF⍺ (**Figure 1C,D,E**; p<0.01, p<0.0001 versus control). GSK2795039 attenuated ROS, nitrite and TNF⍺ levels in a concentration-dependent manner (**Figure 1C,D,E**; p<0.0001 versus L/N). MCC950 reduced ROS but had no effect on nitrite and TNF⍺ levels (Figure 1 C, D, E; p<0.05 vs L/N). These findings suggests that selective NOX2 inhibition disrupts downstream signalling including ROS, nitrite, and TNF⍺, which are key mediators of the microglial pro-inflammatory response.

### NOX2 inhibition attenuates NLRP3 inflammasome activation in pro-inflammatory microglia

To more deeply investigate the NOX2/ROS/NLRP3 inflammatory axis, components of the NLRP3 inflammasome were analysed in cell lysates (CL) and corresponding supernatants (SN) by Western blot. We examined apoptosis-associated speck-like protein containing a CARD (ASC), which is required for the formation of the NLRP3 inflammasome and binding to pro-caspase 1 [42]. Cleaved capsase-1 and cleaved IL-1β were also assessed to measure NLRP3 inflammasome activity. In the CL, L/N modestly increased NLRP3 expression, whereas in the corresponding SN there was a robust increased NLRP3 expression in pro-inflammatory microglia (**Figure 2A**). Although there was no difference in pro-caspase 1 expression, L/N increased cleaved-caspase-1 in the CL in microglia (**Figure 2A**; p<0.05 versus control). Notably, L/N upregulated both pro- and cleaved caspase-1 in the SN. In addition, there was increased pro- and cleaved-IL-1β in both the CL and SN (**Figure 2A**; p<0.0001 vs control). When compared to control, there was no L/N stimulated induction of ASC in the CL, but there was a robust increase in ASC expression in SN (**Figure 2A**).

**Figure 2:**
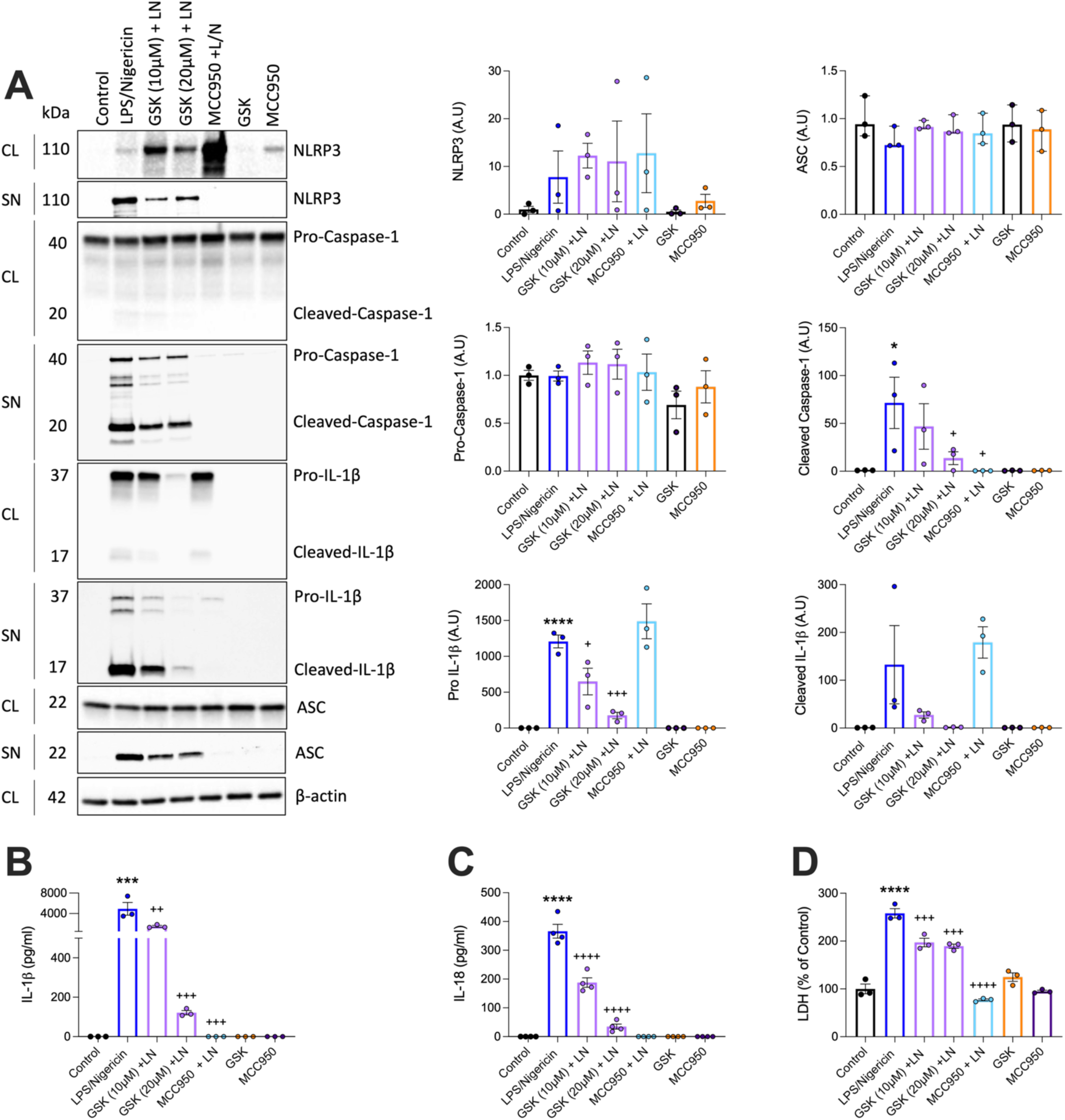
NOX2 inhibition attenuates NLRP3 inflammasome activation in IMGs. IMGs pre-treated with GSK2795039 (10 or 20 µM) or MCC950 (0.5 µM) for 1h prior to stimulation with LPS/nigericin (L/N; 10 µM; 30 min) in a 12-well plate. GSK2795039 (10 µM) and MCC950 (0.01 µM) were used as controls. Cell lysates (CL) and supernatants (SN) were processed and NLRP3, Caspase-1, IL-1β and ASC protein expression were examined. IL-1β, IL-18 and lactate dehydrogenase (LDH) were measured in the conditioned media. In CL, L/N increased NLRP3, cleaved-caspase-1, pro- and cleaved-IL-1β protein expression as shown by immunoblot and corresponding quantification (A; *p<0.05; ****p<0.0001) compared to controls. GSK2795039 reduced cleaved-caspase-1, pro-IL-1β (A; ^+^p<0.05; ^+++^p<0.001) and cleaved-IL-1β but failed to reach significance, versus L/N. MCC950 reduced cleaved-caspase-1 (A; ^+^p<0.05) versus L/N. In the corresponding SN, GSK2795039 reduced L/N induced NLRP3, pro- and cleaved-caspase-1, pro- and cleaved-IL-1β and ASC (A), which were also reduced with MCC950. L/N increased IL-1β, IL-18, and LDH (B-D; ***p<0.001; ****p<0.0001) compared to control. GSK2795039 and MCC950 reduced IL-1β, IL-18, and LDH (B-D; ^++^p<0.01; ^+++^p<0.001; ^++++^p<0.0001) compared to L/N. Data are mean ± SEM (n=3 per group) and are a representative of three independent experiments. *p<0.05; ***p<0.001; ****p<0.0001 versus control, ^+^p<0.05; ^++^p<0.01; ^+++^p<0.001; ^++++^p<0.0001 versus L/N; one-way ANOVA with post hoc Dunnett’s multiple comparisons test versus L/N.

NOX2 inhibition by GSK2795039 had no significant effect on NLRP3 protein expression in CL, but there was a qualitative reduction of NLRP3 expression in SN, as shown by immunoblot (**Figure 2A**). Surprisingly, the NLRP3 inflammasome inhibitor, MCC950, did not reduce NLRP3 expression in the CL in microglia, but its expression was completely abrogated in the corresponding SN (**Figure 2A**). GSK2795039 had no effect on pro-caspase 1 expression in the CL but reduced cleaved-caspase-1 in the CL (Figure 2A; p<0.05 vs L/N). In the corresponding SN, GSK2795039 qualitatively reduced pro- and cleaved-caspase 1 expression in a concentration-dependent manner. MCC950 had no effect on pro-caspase 1 expression in the CL but reduced cleaved-caspase-1 in the CL (**Figure 2A**; p<0.05 vs L/N). Again, in the corresponding SN, MCC950 completely abrogated L/N induced pro- and cleaved-caspase 1 expression. In the CL, GSK2795039 reduced pro-IL-1β in a concentration-dependent manner (**Figure 2A**; p<0.05; p<0.01 vs L/N), as well as cleaved IL-1β, although this failed to reach statistical significance. In the SN, GSK2795039 reduced pro- and cleaved-IL-1β expression in a concentration-dependent manner. MCC950 had no effect on either pro- or cleaved-IL-1β in the CL, but levels of both were reduced in the supernatants. Despite no effect of GSK2795039 on ASC protein expression in the CL, it reduced its expression in the SN. MCC950 did not alter ASC levels in CL but robustly reduced its expression in the SN.

Independent analysis of the SN demonstrated that L/N increased IL-1β and IL-18 (**Figure 2B,C**; p<0.001; p<0.0001 versus control) by ELISA. These NLRP3 inflammasome associated cytokines were reduced by GSK2795039 treatment in a concentration-dependent manner, and as predicted, MCC950 also reduced IL-1β and IL-18 in pro-inflammatory microglia (**Figure 2B,C**; p<0.01; p<0.001; p<0.0001 versus L/N). We next investigated if NOX2 inhibition of IL-1β and IL-18 is pyroptosis dependent. L/N increased lactate dehydrogenase (LDH) (**Figure 2D**; p<0.0001 vs control), an indicator of pyroptotic cell death [43], while GSK2795039 attenuated LDH levels in pro-inflammatory microglia (**Figure 2D**; p<0.05; p<0.0001 vs L/N). As predicted, MCC950 treatment returned LDH to control levels (**Figure 2D**; p<0.0001 vs L/N). These findings indicate that reduced ASC and cleaved caspase 1, by selective NOX2 inhibition, prevents the activation of the NLRP3 inflammasome in pro-inflammatory microglia, which is confirmed by the reduction of downstream products, IL-1β, IL-18 and LDH.

### GSK2795039 attenuates LPS/nigericin ROS, IL-1β, and LDH in primary microglia

We next sought to replicate our findings using primary microglia derived from p1 rat pups. Stimulation of primary microglia with L/N increased ROS, IL-1β, LDH and reduced cell viability (**Figure 3A-D**; p<0.01; p<0.001; p<0.0001 versus control). NOX2 inhibition in primary microglia using GSK2795039 attenuated L/N induced ROS, IL-1β and LDH levels (**Figure 3A-C**; p<0.05; p<0.01; p<0.0001 versus L/N), while it had no effect on cell viability (**Figure 3D**). The NLRP3 inhibitor MCC950 had no effect on ROS and IL-1β (**Figure 3A,B**), albeit mainly due to data scatter and variability. MCC950 reduced LDH (**Figure 3C**; p<0.05 versus L/N) but did not rescue L/N induced-cell death (**Figure 3D**). These findings in primary microglia confirmed prior studies in IMG microglia and demonstrate NOX2 inhibition by GSK2795039 attenuates L/N elicited ROS production and NLRP3 inflammasome activation.

**Figure 3:**
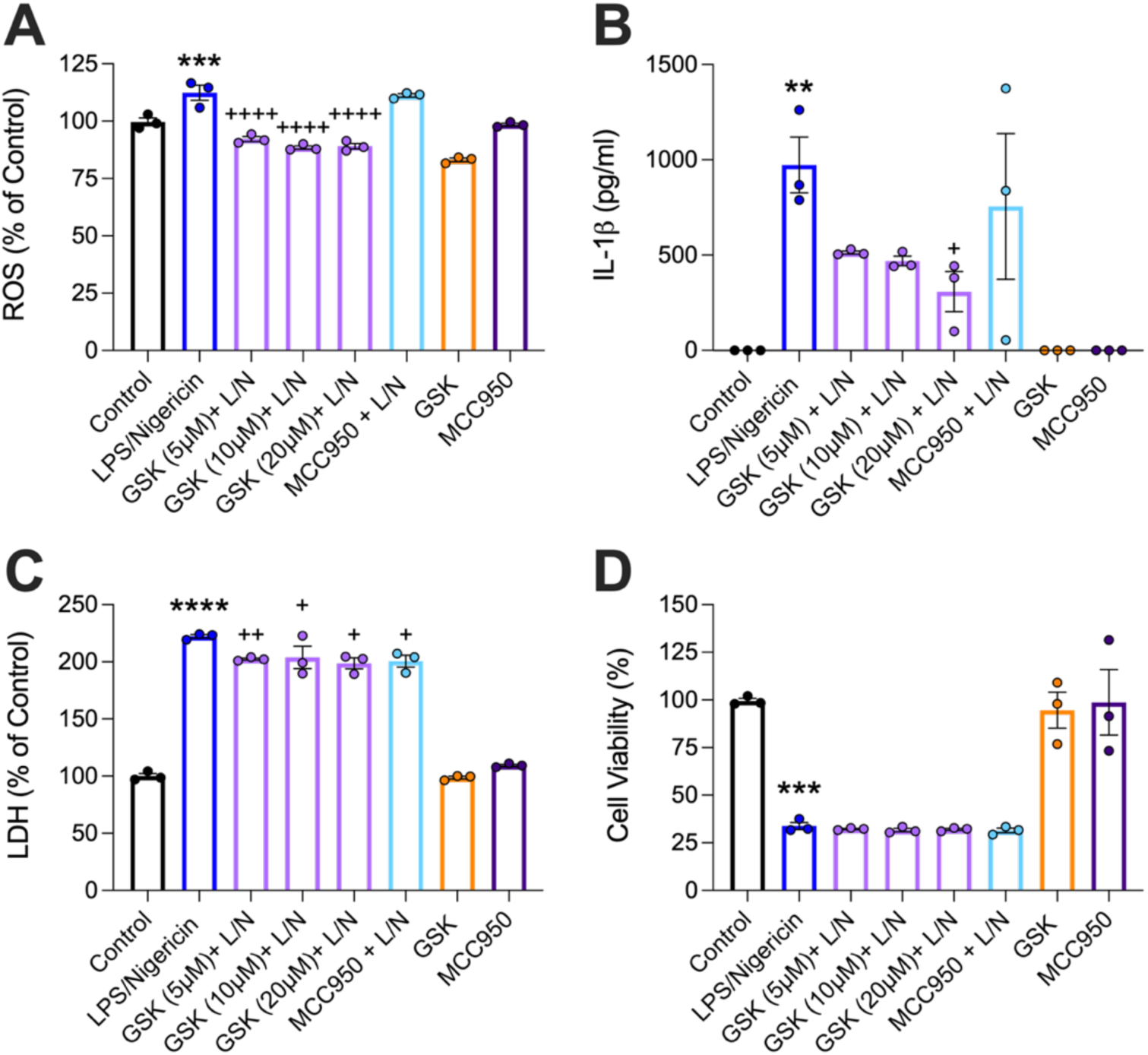
GSK2795039 attenuated L/N induced NOX2 and NLRP3 inflammasome activation in primary microglia. Primary microglia were pre-treated with GSK2795039 (5, 10, or 20 µM) or MCC950 (0.01 µM) for 1h prior to stimulation with LPS/nigericin (L/N; 10 µM; 30 min) in a 96-well plate. GSK2795039 (10 µM) and MCC950 (0.01 µM) were used as controls. ROS and cell viability were measured in the cells, LDH and IL-1β were measured in the conditioned media. L/N increased ROS, IL-1β, LDH and reduced cell viability (A-D; **p<0.01; ***p<0.001; ****p<0.0001) compared to control. GSK2795039 reduced ROS, IL-1β and LDH (A-C; ^+^p<0.05; ^++^p<0.01; ^++++^p<0.0001) versus L/N. MCC950 reduced LDH (D; ^+^p<0.05) compared to L/N. GSK2795039 or MCC950 did not rescue cell viability loss (D). Data are mean ± SEM (n=3 per group) and are a representative of three independent experiments. **p<0.01; ***p<0.001; ****p<0.0001 versus control, ^+^p<0.05, ^++^p<0.01, ^++++^p<0.0001 versus L/N; one-way ANOVA with post hoc Dunnett’s multiple comparisons test versus L/N.

### Cellular players in NOX2-ROS-NLRP3 inflammatory axis activation in the injured brain

NOX2 activation in microglia/macrophages contributes to post-traumatic neuroinflammation and associated neurodegeneration up to 1-year post-TBI [2], with pharmacological or genetic NOX2 inhibition resulting in reduced neuroinflammation and improved neurological recovery in injured mice [4, 44]. In addition, inhibition of NLRP3 inflammasome activation using MCC950 reduces post-traumatic neuroinflammation in mice [26, 27, 45]. Together, these findings indicate that both NOX2 and NLRP3 inflammasome activation contribute to progressive neuroinflammation following TBI, but the cellular contributors to inflammatory pathway activation are poorly defined.

We therefore used flow cytometry to examine cellular profiles of the NOX2-ROS-NLRP3 inflammatory axis in a controlled cortical impact (CCI) of experimental TBI in mice. Adult male C57Bl6J mice were subjected to sham or moderate-level CCI [30] and cohorts of mice were humanely euthanized at 1-, 3- and 7-days post-injury (DPI) for cellular and molecular analysis. NOX2 related, *Cyba*, *Cybb*, the inflammasome, *Nlrp3*, and microglial phagocytic, *Cd68*, genes were robustly increased in the ipsilateral hippocampus following TBI with expression peaking at 3 DPI (**Figure 4A**; p<0.05 vs sham). *Cd68* was significantly reduced at 7 DPI (**Figure 4A**; p<0.05 vs 3 DPI), with a reduction in *Cyba*, *Cybb*, and *Nlrp3*, which failed to reach statistical significance. Pro-inflammatory genes were also increased in the hippocampus with a significant upregulation of *Il1b* and *Tnf* mRNAs that peaked at 1 DPI (**Figure 4A**; p<0.001 vs sham), with reduced expression of both genes at 3 and 7 DPI (**Figure 4A**; p<0.05; p<0.01; p<0.001 vs 1 DPI).

**Figure 4:**
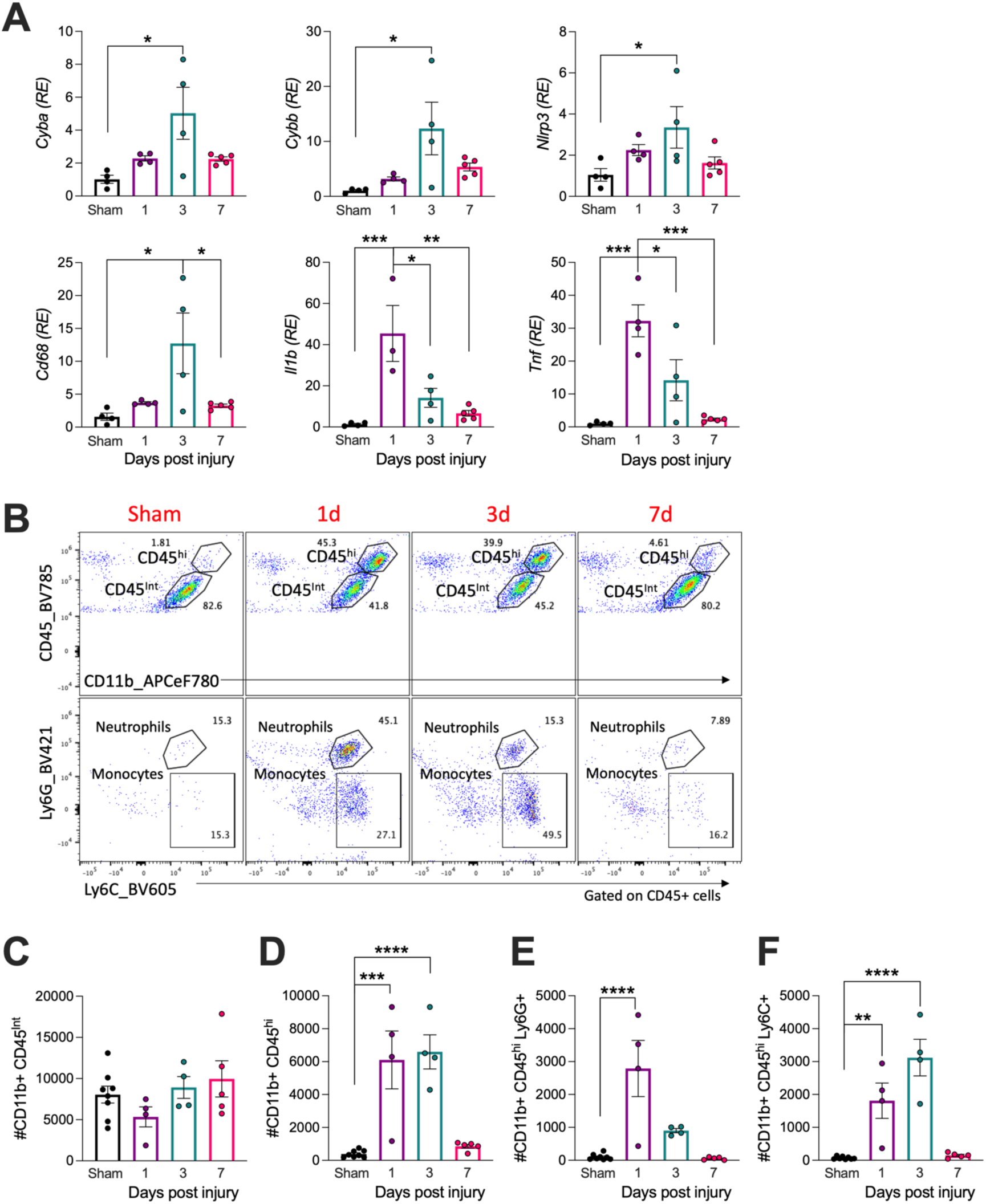
NOX2 and NLRP3 mRNA expression upregulated at 3 DPI and increased infiltration of neutrophils and monocytes at 1-3 DPI. Adult male C57BL6/J mice were subjected to sham or CCI surgery and euthanised at 1-, 3- or 7 DPI. Tissue punches from the ipsilateral hippocampus of sham or CCI mice were collected for gene expression analysis. CCI induced increased *Cyba*, *Cybb*, *Cd68* and *Nlrp3* mRNA at 3 DPI (A; *p<0.05 vs sham) and there was significantly lower *Cd68* mRNA at 7DPI (A; *p<0.05 vs 3 DPI). CCI increased *Il1b* and *Tnf* gene expression at 1 DPI (A; ***p<0.001 vs sham), which was lower at 3 and 7 DPI (A; *p<0.05, **p<0.01, ***p<0.001 vs 1 DPI). CCI induced increased number of infiltrating myeloid cells (CD11b^+^CD45^hi^) at 1- and 3 DPI as shown in the representative dot plots (B) and quantification (D; ***p<0.001; ****p<0.0001 vs sham). The number of neutrophils (Ly6G^+^) peaked at 1 DPI (B,E; ****p<0.0001 vs sham), and the number of monocytes (Ly6C^+^) peaked at 3 DPI (B,F; **p<0.01; ****p<0.0001 vs sham). Data are mean ± SEM (n=4-5 per group). *p<0.05; **p<0.01; ***p<0.001; ****p<0.0001; by One-Way ANOVA with Tukey’s multiple comparisons test.

We next sought to determine cellular changes in the injured brain. The number and phenotype of brain resident microglia and infiltrating innate immune cells were evaluated at 1, 3 and 7 DPI by multi-dimensional flow cytometry analysis. When compared to sham levels, there were no changes in microglial numbers (CD11b^+^CD45^int^) after TBI (**Figure 4B,C**), while there was increased infiltration of peripherally derived myeloid cells (CD11b^+^CD45^hi^) into the brain at 1 and 3 DPI (**Figure 4B,D**; p<0.001; p<0.0001 vs sham). Deeper analysis demonstrated that neutrophils (CD11b^+^CD45^hi^Ly6G^+^) rapidly accumulated early after TBI and were the predominant population at 1 DPI (**Figure 4B,E**; p<0.0001 vs sham). Monocytes (CD11b^+^CD45^hi^Ly6C^+^) also accumulated in the injured brain, with peak numbers appearing at 3 DPI (**Figure 4B,F**; p<0.01; p<0.0001 vs sham).

We then examined functional changes in immune cells in the injured brain. Microglia were first investigated. When compared to sham, TBI increased NOX2^+^ microglia at 3 and 7 DPI (**Figure 5A,B**; p<0.0001 vs sham). Although non-significant, there were increased IL-1β^+^ microglia at 3 DPI (**Figure 5A,C**). TBI upregulated double positive (DP) NOX2^+^ IL-1β^+^ at 3 and 7 DPI (**Figure 5A,D**; p<0.05; p<0.001 vs sham; see Q2 in representative dot plots for DP populations in **Figure 5A**). TBI also robustly increased microglial ROS production across all timepoints (**Figure 5M,N**; p<0.05; p<0.001; p<0.0001 vs sham), with peak microglial ROS production at 3 DPI. When examining neutrophil phenotypes, there were increased NOX2^+^ neutrophils infiltrating the injured brain at 1 and 3 DPI (**Figure 5E,F**; p<0.05; p<0.001 vs sham). There was a significant reduction in NOX2^+^ neutrophils at 7 DPI (**Figure 5E,F**; p<0.05 vs 3 DPI), which was due to an overall decreased number of neutrophils at this timepoint. TBI also increased IL-1β^+^ neutrophils (**Figure 5E,G**; p<0.01 vs sham) and DP NOX2^+^ IL-1β^+^ neutrophils (**Figure 5E,H**; p<0.01 vs sham; see Q2 in representative dot plots for DP populations in **Figure 5E**) at 1 DPI. TBI also increased neutrophil ROS production at 1 DPI (**Figure 5M,O**; p<0.05 vs sham). Finally, monocyte phenotypes were examined, and there were increased NOX2^+^ monocytes infiltrating the injured brain at 1 and 3 DPI (**Figure 5I,J**; p<0.001; p<0.0001 vs sham). There was a significant reduction in NOX2^+^ monocytes at 7 DPI (**Figure 5I,J**; p<0.001 vs 3 DPI), which was due to an overall decreased number of monocytes at this timepoint. TBI greatly increased IL-1β^+^ monocytes (**Figure 5I,K** p<0.0001 vs sham) and DP NOX2^+^ IL-1β^+^ monocytes (**Figure 5I,L**; p<0.0001 vs sham; see Q2 in representative dot plots for DP populations in **Figure 5I**) at 1 DPI. TBI also increased monocytic ROS production at 1 and 3 DPI (**Figure 5M,P**; p<0.05; p<0.0001 vs sham), with peak monocyte ROS production at 3 DPI. These comparative studies allow for insights into NOX2/ROS/IL-1β functional interactions in different cellular populations in the injured brain.

**Figure 5:**
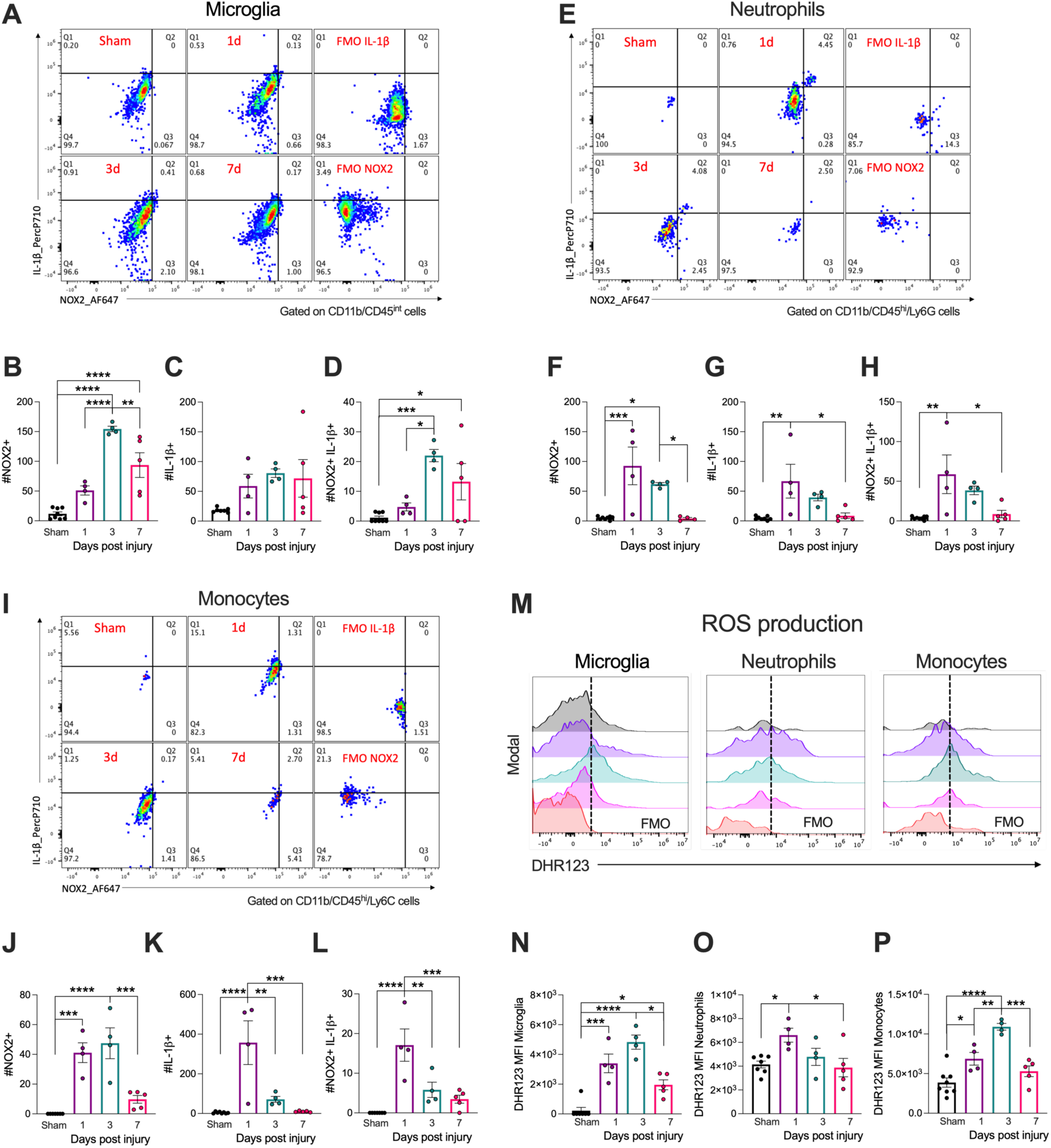
Microglia, neutrophil, and monocyte NOX2/ROS/IL-1β production peaked at 1- and 3 DPI. Adult male C57BL6/J mice were subjected to sham or CCI surgery and euthanised at 1-, 3- or 7 DPI. Mononuclear cells isolated from the ipsilateral cortex were stained with surface markers, DHR123 (ROS dye), followed by intracellular staining of NOX2 and IL-1β and analysed by flow cytometry. CCI increased #NOX2^+^ and #NOX2^+^IL- 1β^+^ microglia at 3- and 7 DPI as shown in the representative quadrants (A) and corresponding quantifications (B, D; *p<0.05; ***p<0.001; ****p<0.0001 vs sham). CCI increased #NOX2^+^, #IL-1β^+^, and #NOX2^+^IL-1β^+^ neutrophils and monocytes at 1 DPI as shown in the representative quadrants (E, I) and corresponding quantifications (F-H, J-L; **p<0.01; ***p<0.001; ****p<0.0001 vs sham). Numbers reduced to sham levels at 7 DPI (F-H, J-L; *p<0.05; ***p<0.001 vs 1 and 3 DPI). Monocytic #NOX2^+^ peaked at 3 DPI (I, J; ****p<0.0001 vs sham). CCI induced microglial ROS production across all time-points (M, N; *p<0.05; ***p<0.001; ****p<0.0001 vs sham), which peaked at 3 DPI (N; *p<0.05 vs 7 DPI). Neutrophils and monocytes induced ROS production at 1 DPI, (M, O, P; *p<0.05 vs sham), which was reduced at 7 DPI (O, P; *p<0.05; ***p<0.001 vs 1 and 3 DPI). Monocytic ROS peaked at 3 DPI (M, P; **p<0.01 vs 1 DPI), and was reduced at 7 DPI (P; ***p<0.001 vs 3 DPI). Data are mean ± SEM (n=4-5 per group). *p<0.05; **p<0.01; ***p<0.001; ****p<0.0001; by One-Way ANOVA with Tukey’s multiple comparisons test.

### The small molecule NOX2 inhibitor, GSK2795039, has good brain penetration and modestly improves neurological function in the acute phase after TBI

We then sought to determine whether GSK2795039 could alter post-traumatic neuroinflammation and neurological outcomes after experimental TBI. The small molecule NOX2 inhibitor, GSK2795039, reduces inflammation *in vivo* in mice [25], and we wanted to check its brain distribution levels after TBI, so we performed *in vivo* pharmacokinetics (PK) analysis. Adult C57BL6/J male mice were subjected to CCI and GSK2795039 (100mg/kg) or Vehicle was administered intraperitoneally (i.p.) at 2h post-injury. Mice were humanely euthanized 0.5h later, and blood, ipsilateral cortex and hippocampus, and liver was collected for PK analysis (**Figure 6Ai**). The measured GSK2795039 levels in the cortex (8.19% of plasma) and hippocampus (7.01% of plasma; **Figure 6Aii**) are consistent with literature of 5-10% of plasma levels needed for drug penetration in brain [46]. As expected, GSK2795039 was also detected in the liver (37.09% of plasma).

**Figure 6:**
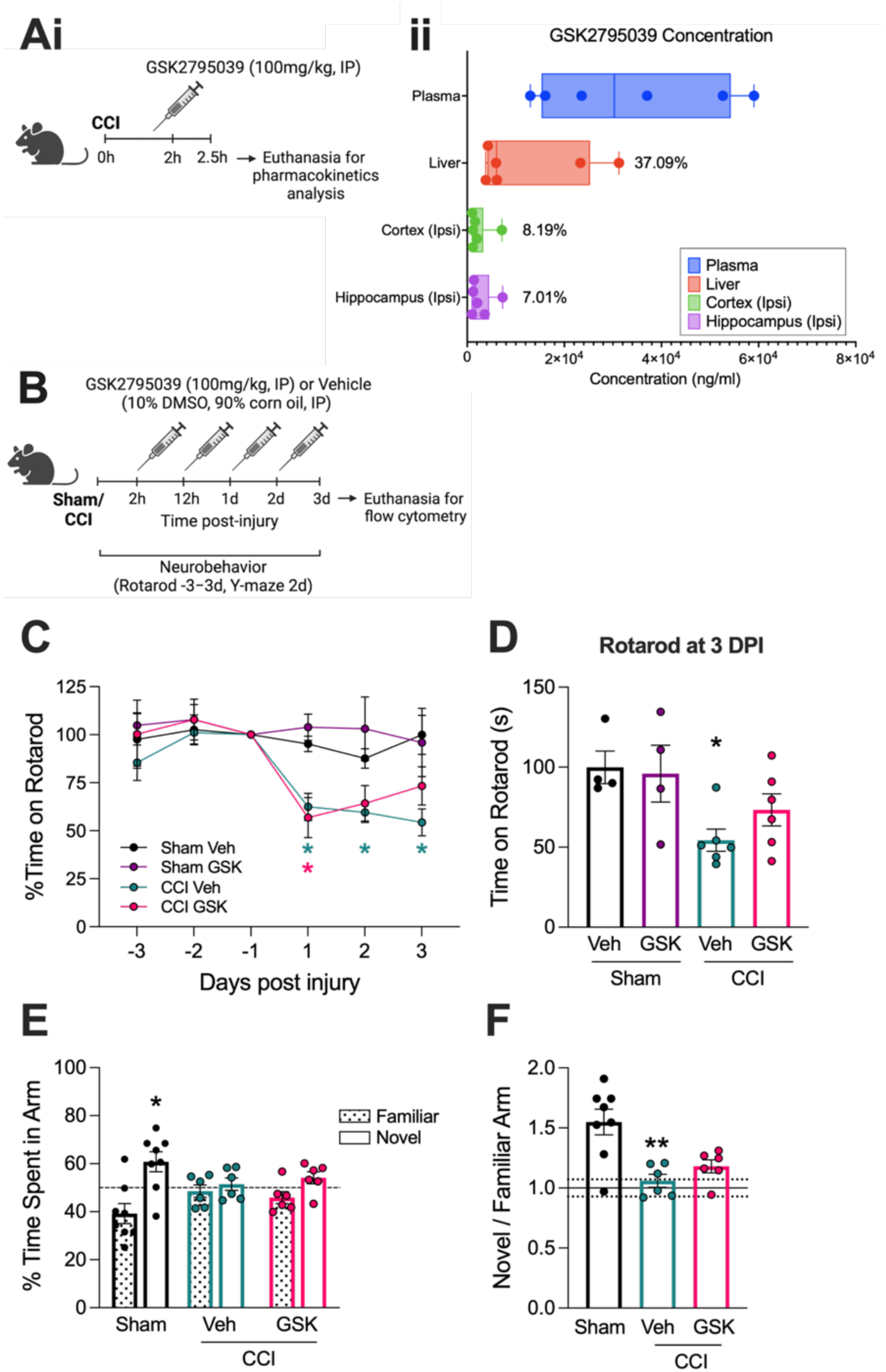
GSK2795039 has good brain penetration and modestly improves neurological function in the acute phase after TBI. GSK2795039 (100mg/kg; i.p.) was administered 2h following CCI in adult male C57BL6/J mice, and mice were sacrificed 30 min later (Ai). The concentration of GSK2795039 (ng/ml) in plasma, liver, cortex and hippocampus was quantified using LC–MS analysis resulting in GSK2795039 levels in liver (37.09% of plasma), cortex (8.19%) and hippocampus (7.01%) (Aii). In a separate cohort of mice, GSK2795039 (100mg/kg; i.p.) or vehicle was administered systemically starting at 2h post-CCI (B). Gross motor function was measured by an accelerating rotarod at 1-,2- and 3 DPI (B). Vehicle treated CCI mice spent reduced time on the rotarod at 1-, 2- and 3 DPI (C; *p<0.05 vs sham veh). GSK2795039 treated CCI mice spent increased time on the rotarod at 3 DPI, but levels failed to reach significance (C, D). Spatial learning and memory were assessed by a two-trials Y-Maze at 2 DPI (B). Sham mice spent more time in the novel arm of the Y-Maze (E; *p<0.05) compared to the familiar arm, while vehicle treated CCI mice did not show any difference in the time spent in the familiar vs novel arm. GSK2795039 treated CCI mice spent more time in the novel arm, but levels failed to reach statistical significance (E). The fold difference of the novel arm (over the familiar arm) was significantly lower in vehicle treated CCI mice (F; **p<0.01 vs sham). GSK2795039 treated CCI mice showed a trend towards sham levels, but this failed to reach statistical significance. Dotted lines represent mean ± SEM of the time spent in the familiar arm. Data are mean ± SEM (n=4-6 per group). *p<0.05 vs sham veh/sham GSK2795039; **p<0.01 vs sham; Two-Way ANOVA with repeated measures and uncorrected Fisher’s LSD (C, D); paired t-test (E); One-Way ANOVA with Dunnett’s post hoc test (F).

We next investigated whether GSK2795039 treatment starting at 2h post-injury could attenuate NOX2/NLRP3 inflammasome activation and improve neurological recovery acutely after TBI. Adult male C57Bl6/J mice were subjected to sham or CCI and administered GSK2795039 (100mg/kg; i.p.) at 2h, 12h, 1d, and 2d post-injury (**Figure 6B**). Gross motor function was assessed using an accelerating rotarod at 1, 2 and 3 DPI. As expected, vehicle treated CCI mice spent reduced time on the rotarod at 1, 2 and 3 DPI (**Figure 6C**; p<0.05 vs sham veh). GSK2795039 treated CCI mice spent reduced time on the rotarod at 1 DPI (**Figure 6C**; p<0.05 vs sham GSK2795039), but showed improvements at 2 and 3 DPI, with levels not statistically different to sham (**Figure 6C**). Sub analysis of rotarod performance at 3 DPI demonstrated that vehicle treated CCI mice spent 54.3 ± 6.9 sec on the rotarod, whereas GSK2795039 treated CCI mice spent 73.3 ± 10.0 sec on the rotarod (**Figure 6D**; Sham + Veh = 99.9 ± 10.2 sec; Sham + GSK2795039 = 95.9 ± 17.2 sec).

We used a two-trial Y-maze test to assess spatial memory in sham and injured mice at 2 DPI. Sham mice (Vehicle and GSK2795039 groups pooled) spent more time in the novel arm compared to the familiar arm (**Figure 6E**; p<0.05 vs sham familiar). Vehicle treated CCI mice did not differentiate the novel from the familiar arm (**Figure 6E**), with only half of the mice spending more than 50% (chance) in the novel arm. In contrast, GSK2795039 treated CCI mice spent more time in the novel arm (**Figure 6E**), with 5 out of the 6 mice spending more than 50% of the time in the novel arm (chance). The ratio of novel over the familiar arm was also assessed. Vehicle treated CCI mice had a decreased ratio (**Figure 6F**; p<0.01 vs sham), whereas GSK2795039 treated CCI mice had an increased ratio, but levels failed to reach statistical significance. There was no difference in distance travelled in the Y-maze test between the groups (Sham = 12.31 ± 0.7895m; + CCI Veh = (13.69 ± 1.411m); CCI GSK2795039 + = 11.80 ± 0.9463m).

### NOX2 inhibition reduces NOX2^+^ IL-1β^+^ infiltrating myeloid cells in the injured brain

We then used flow cytometry to investigate the effects of NOX2 inhibition on the inflammatory phenotype of microglia and infiltrating myeloid cells after TBI. When compared to vehicle treated CCI mice, GSK2795039 treated CCI mice had reduced numbers of infiltrating monocytes at 3DPI, albeit no difference in infiltrating neutrophil or microglial numbers were observed (**Table 1**). As expected, vehicle treated CCI mice had increased numbers of NOX2^+^ and IL-1β^+^ microglia (CD11b^+^CD45^Int^) and infiltrating myeloid cells (CD11b^+^CD45^hi^) (**Figure 7A-F**; p<0.0001 vs sham veh). GSK2795039 treated CCI mice had reduced numbers of NOX2^+^ and IL-1β^+^ microglia, but levels failed to reach statistical significance (**Figure 7A-D**). Notably, GSK2795039 treated CCI mice had reduced numbers of NOX2^+^ and IL-1β^+^ infiltrating myeloid cells (**Figure 7A,B,E,F**; p<0.001 versus CCI Veh).

**Figure 7:**
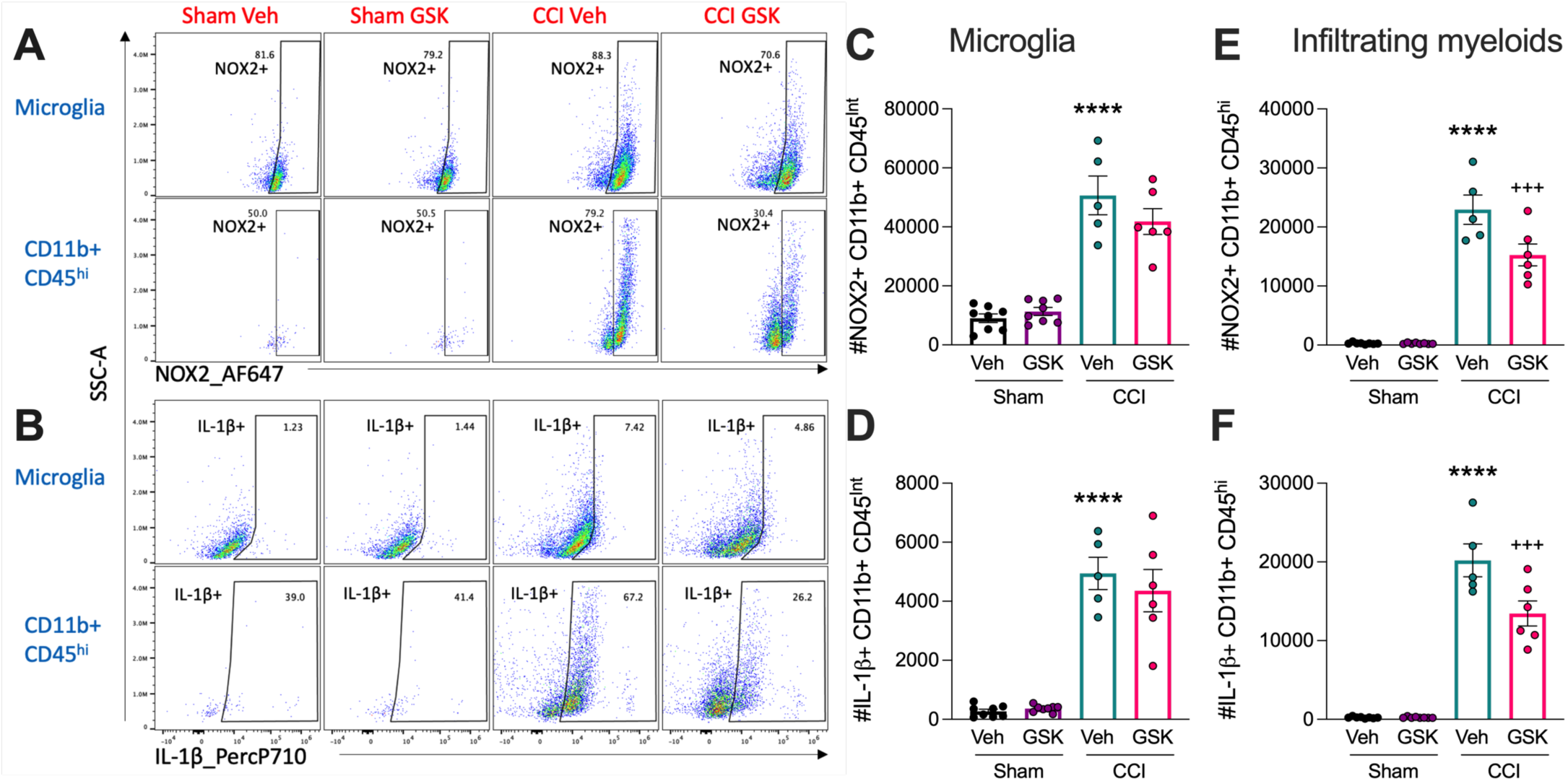
Pharmacological NOX2 inhibition reduces NOX2^+^ IL-1β^+^ infiltrating myeloid cells in the injured brain. GSK2795039 was administered at 2h, 12h, 1d and 2d post CCI. Mononuclear cells isolated from the ipsilateral cortex were stained with surface markers followed by intracellular staining of NOX2 and IL-1β and analysed by flow cytometry. CCI increased the absolute number of NOX2^+^ and IL-1β^+^ CD11b^+^CD45^Int^ microglia and CD11b^+^CD45^hi^ infiltrating cells in vehicle treated CCI mice (C-F; ****p<0.0001 vs sham veh). GSK2795039 treated CCI mice had reduced #NOX2^+^ #IL-1β^+^ microglia, but levels failed to reach significance (C). GSK2795039 treated CCI mice had significantly reduced #NOX2^+^ IL-1β^+^ infiltrating cells (E, F; ^+++^p<0.001 vs CCI veh). Data are mean ± SEM (n=4-6 per group). ****p<0.0001 vs sham veh, ^+++^p<0.001 vs CCI veh; Two-Way ANOVA with uncorrected Fisher’s LSD.

**Table 1:**
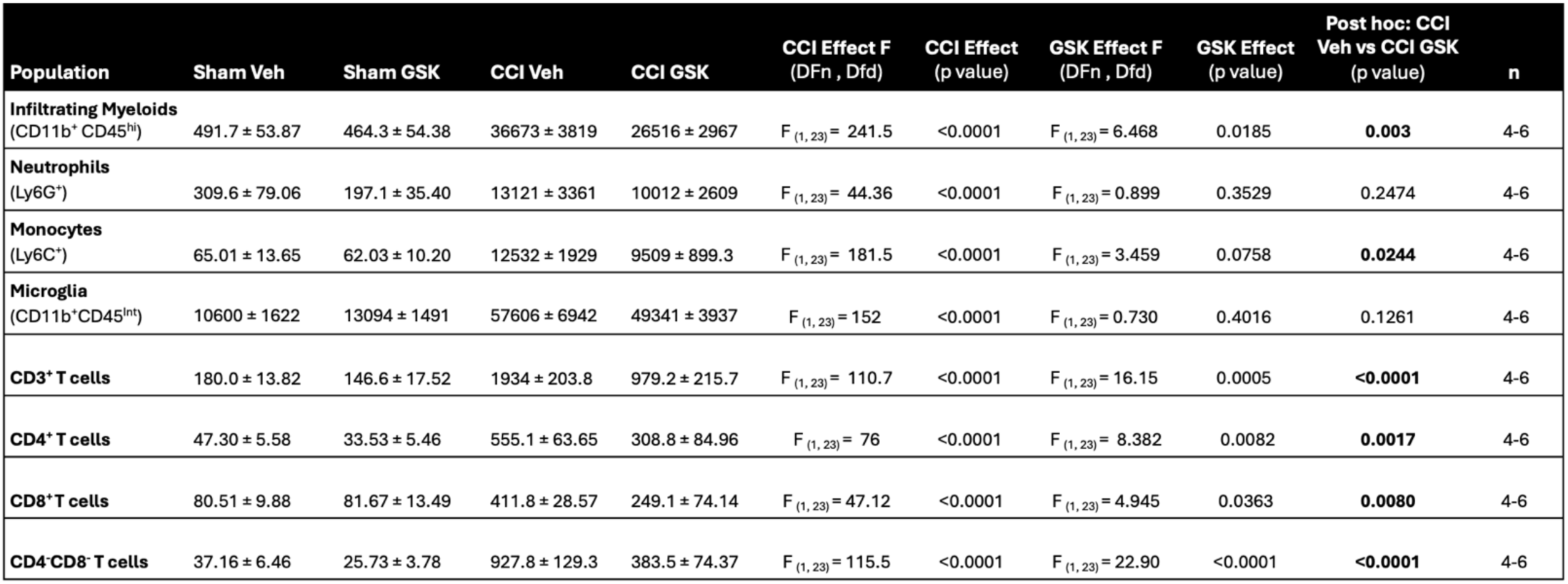
The effect of CCI and GSK2795039 treatment on immune cell population numbers in the brain at 3 DPI. Abbreviations: CCI, controlled cortical impact; Veh, Vehicle; GSK, GSK2795039.

We then performed a deeper analysis on neutrophil and monocyte populations in the injured brain. Vehicle treated CCI mice had increased numbers of NOX2^+^ and IL-1β^+^ neutrophils (CD11b^+^CD45^hi^Ly6G^+^) as well as DP NOX2^+^IL-1β^+^ neutrophils (**Figure 8Ai-iv**; p<0.0001 vs sham veh). Notably, GSK2795039 treated CCI mice had reduced numbers of IL-1β^+^, NOX2^+^, and DP NOX2^+^IL-1β^+^ neutrophils (**Figure 8Ai-iv**; p<0.05; p<0.01; p<0.0001 vs CCI veh). When examining monocytes (CD11b^+^CD45^hi^Ly6C^+^), vehicle treated CCI mice had increased numbers of NOX2^+^ and IL-1β^+^ monocytes as well as DP NOX2^+^IL-1β^+^ monocytes (**Figure 8Bi-iv**; p<0.0001 vs sham veh). Notably, GSK2795039 treated CCI mice had reduced numbers of NOX2^+^ monocytes (**Figure 8Bi-iv**; p<0.05 vs CCI veh), and reduced numbers of IL-1β^+^ and DP NOX2^+^IL-1β^+^ monocytes, but levels failed to reach statistical significance. The immunophenotyping findings indicate that NOX2 inhibition attenuates TBI neuroinflammation, with specific effects on NOX2^+^ IL-1β^+^ neutrophils and monocytes that infiltrate the injured brain.

**Figure 8:**
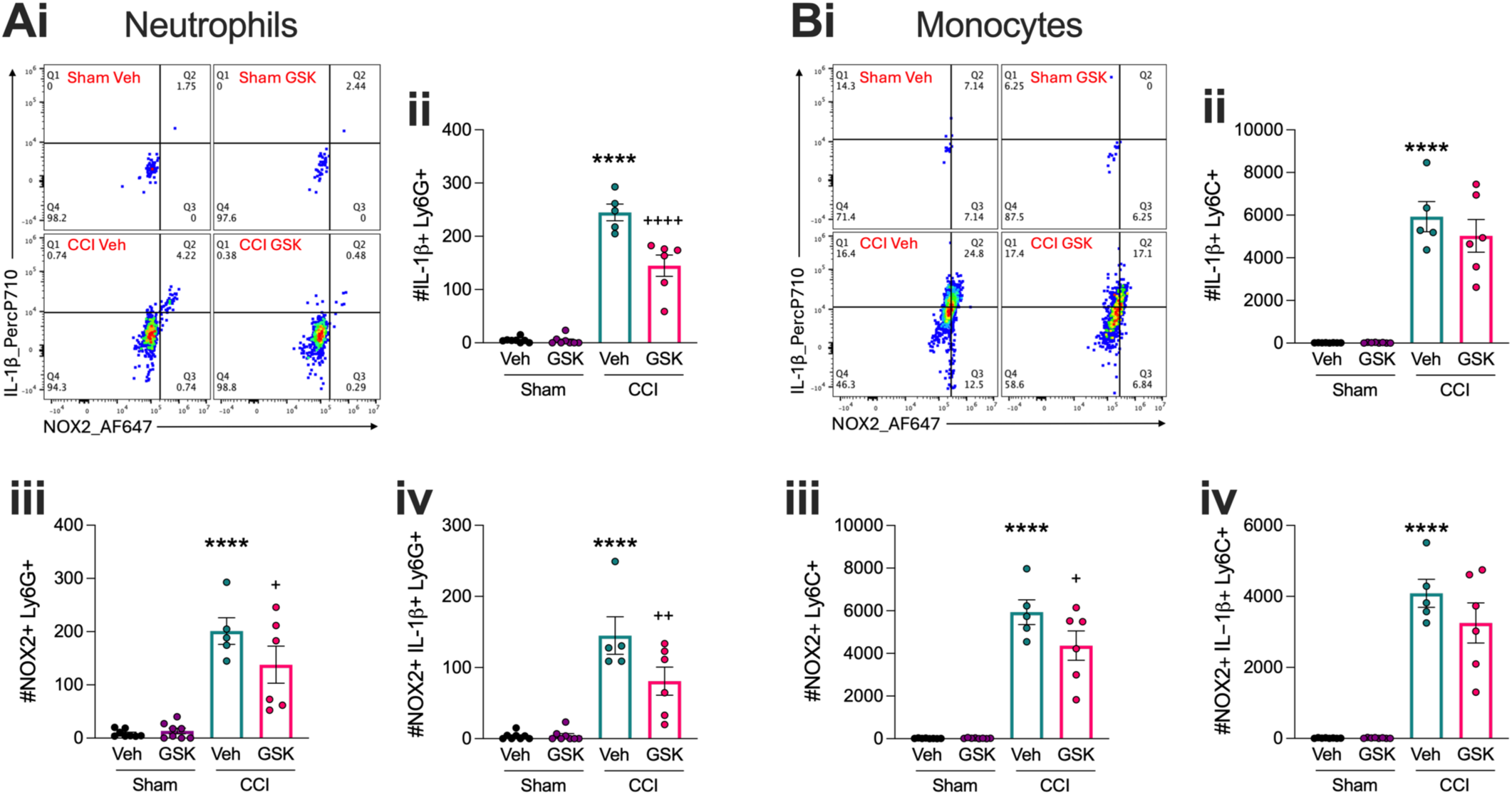
Pharmacological NOX2 inhibition reduces absolute numbers of NOX2^+^ and IL-1β^+^ neutrophils and NOX2^+^ monocytes at 3 DPI. GSK2795039 was administered at 2h, 12h, 1d and 2d post CCI. Mononuclear cells isolated from the ipsilateral cortex were stained with surface markers, followed by intracellular staining of NOX2 and IL-1β, and analysed by flow cytometry. CCI increased the absolute number of NOX2^+^ and IL-1β^+^ neutrophils and monocytes in vehicle treated CCI mice (Ai-iv, Bi-iv; ****p<0.001 vs sham veh). GSK2795039 reduced #NOX2^+^ #IL-1β^+^ neutrophils (Ai-iv; ^+^p<0.05; ^++^p<0.01; ^++++^p<0.0001 vs CCI veh) and #NOX2^+^ monocytes (Biii; ^+^p<0.05 vs CCI veh). Data are mean ±SEM (n=4-6 per group). ****p<0.0001 vs sham veh, ^+^p<0.05; ^++^p<0.01; ^++++^p<0.0001 vs CCI veh; Two-Way ANOVA with uncorrected Fisher’s LSD.

### NOX2 inhibition reduces IL-1R expression on T cells in the injured brain

We next investigated whether NOX2 inhibition could also alter the adaptive immune response following TBI. When compared to vehicle treated CCI mice, GSK2795039 treated CCI mice had reduced numbers of infiltrating CD3^+^ lymphocytes, with significantly reduced numbers of CD4^+^, CD8^+^ and CD4^-^/CD^-^ T cells in the injured brain (**Table 1**). We hypothesised that IL-1R expressing T cells respond to IL-1β released from cells of myeloid origin, such as microglia or monocytes [47]. This ligand binding complex triggers and facilitates T cell responses [48], and thus signalling may act as an inducer of adaptive immune responses in the injured brain. To evaluate this, IL-1R^+^ expression was first gated from total CD3^+^ T cells (**Figure 9A**; upper panel). Of the IL-1R^+^CD3^+^T cells, the number of CD4^+^, CD8^+^ and double negative (DN) CD4^-^/CD8^-^were determined (**Figure 9B**; lower panel). When compared to sham, vehicle treated CCI mice had increased IL-1R expression on CD3^+^, CD4^+^, CD8^+^ and CD4^-^/CD8^-^ T cells (**Figure 9A-E**; p<0.0001 vs sham veh). Notably, GSK2795039 treated CCI mice had reduced numbers of IL-1R^+^CD3^+^, IL-1R^+^CD4^+^, IL-1R^+^CD8^+^ and IL-1R^+^CD4^-^/CD8^-^T cells (**Figure 9A-E**; p<0.05; p<0.01; p<0.0001 vs CCI veh). The reduction in IL-1R^+^ T cells during the acute phase post-injury suggests that myeloid-T cell crosstalk may be altered by NOX2 inhibition in the injured brain.

**Figure 9:**
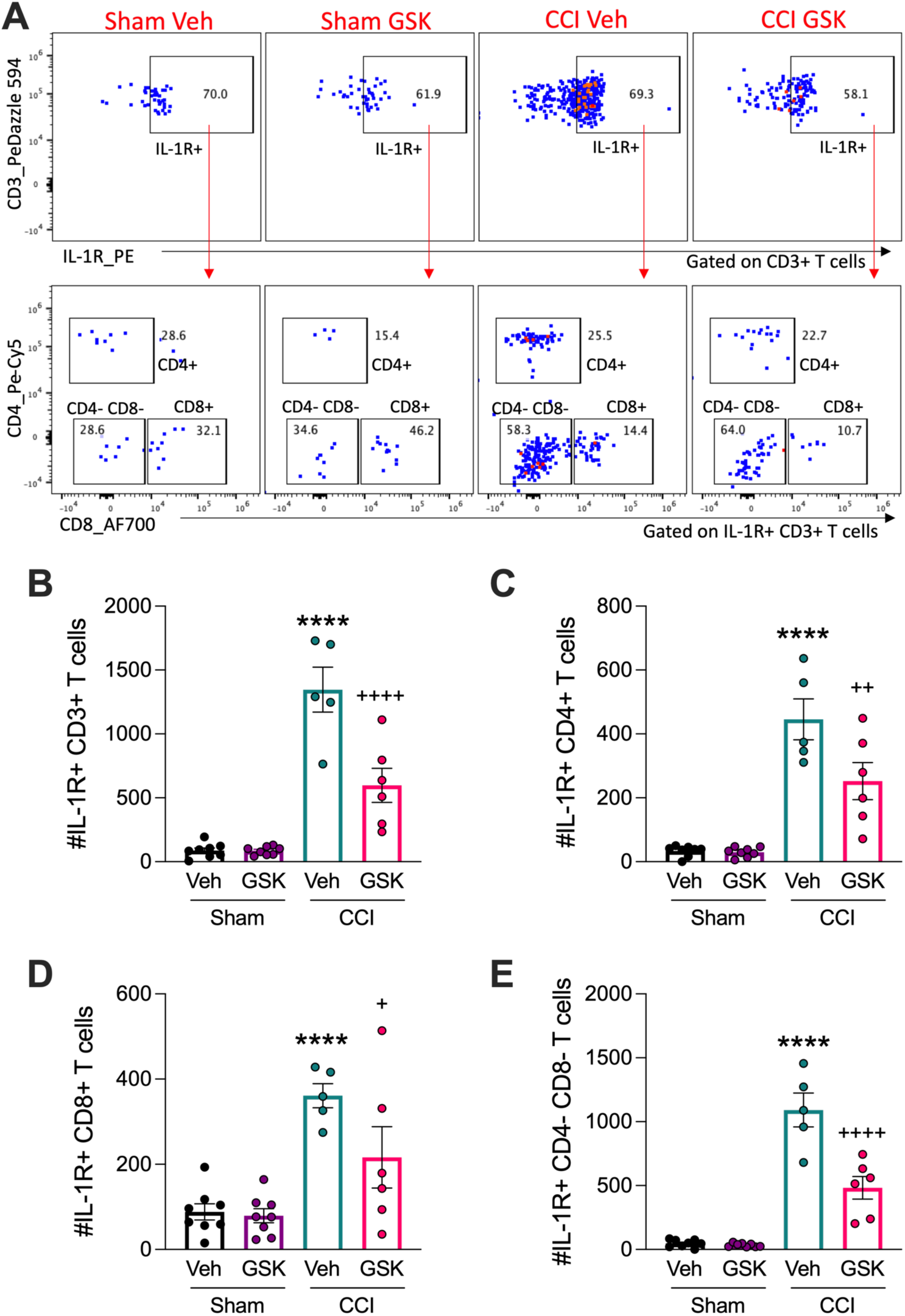
Pharmacological NOX2 inhibition reduces absolute numbers of IL-1R+ T cells at 3 DPI. GSK2795039 was administered at 2h, 12h, 1d and 2d post CCI. Mononuclear cells isolated from the ipsilateral cortex were stained for CD3, CD4, CD8 and IL-1R, and analysed by flow cytometry. CCI increased absolute numbers of IL-1R^+^ expression on CD3^+^, CD4^+^, CD8^+^ and CD4^-^CD8^-^T cells in vehicle treated CCI mice as shown by representative dot plots (A) and corresponding quantifications (B-E; ****p<0.0001 vs sham veh). GSK2795039 treated CCI mice significantly reduced absolute numbers of IL-1R^+^CD3^+^, IL-1R^+^CD4^+^, IL-1R^+^CD8^+^ and IL-1R^+^CD4^-^ CD8^-^T cells (B-E; ^+^p<0.05; ^++^p<0.01; ^++++^p<0.0001 vs CCI veh). Data are mean ±SEM (n=4-6 per group). ****p<0.0001 versus sham, ^+^p<0.05; ^++^p<0.01; ^++++^p<0.0001 vs CCI veh; Two-Way ANOVA with uncorrected Fisher’s LSD.

### GSK2795039 treatment fails to improve long-term neurological recovery following TBI but reduces neuropathology

Finally, we examined the effect of GSK2795039 treatment on chronic neurological and histopathological outcomes following experimental TBI. Adult male C57Bl6/J mice were subjected to sham or CCI and administered GSK2795039 (100mg/kg; i.p.) at 2h, 12h, 1d, 2d, 3d and 7d post-injury. Fine motor coordination was assessed using a beam walk test at baseline prior to sham/CCI and up to 28 DPI, while cognitive function was assessed using the novel object recognition (NOR) test at 17-20 DPI (**Figure 10A**). When compared to vehicle treated sham controls, vehicle treated CCI mice had increased foot-faults on the beam walk test from 1-28 DPI, which resulted in gradual improvements over time (**Figure 10B**; p<0.01; p<0.001; p<0.0001 vs sham). GSK2795039 treated CCI mice also had increased foot-faults from 1-28 DPI (**Figure 10B**; p<0.01; p<0.0001 vs sham). Although there were minor improvements in foot-faults in GSK2795039 treated CCI mice at various timepoints, levels did not reach statistical significance. In the NOR there was a reduced discrimination index (DI) in vehicle treated CCI mice compared to sham, indicative of worsened cognitive function, but levels failed to reach statistical significance (**Figure 10C**). GSK2795039 treated CCI mice had improved DI in this test, but these effects did not reach statistical significance.

**Figure 10:**
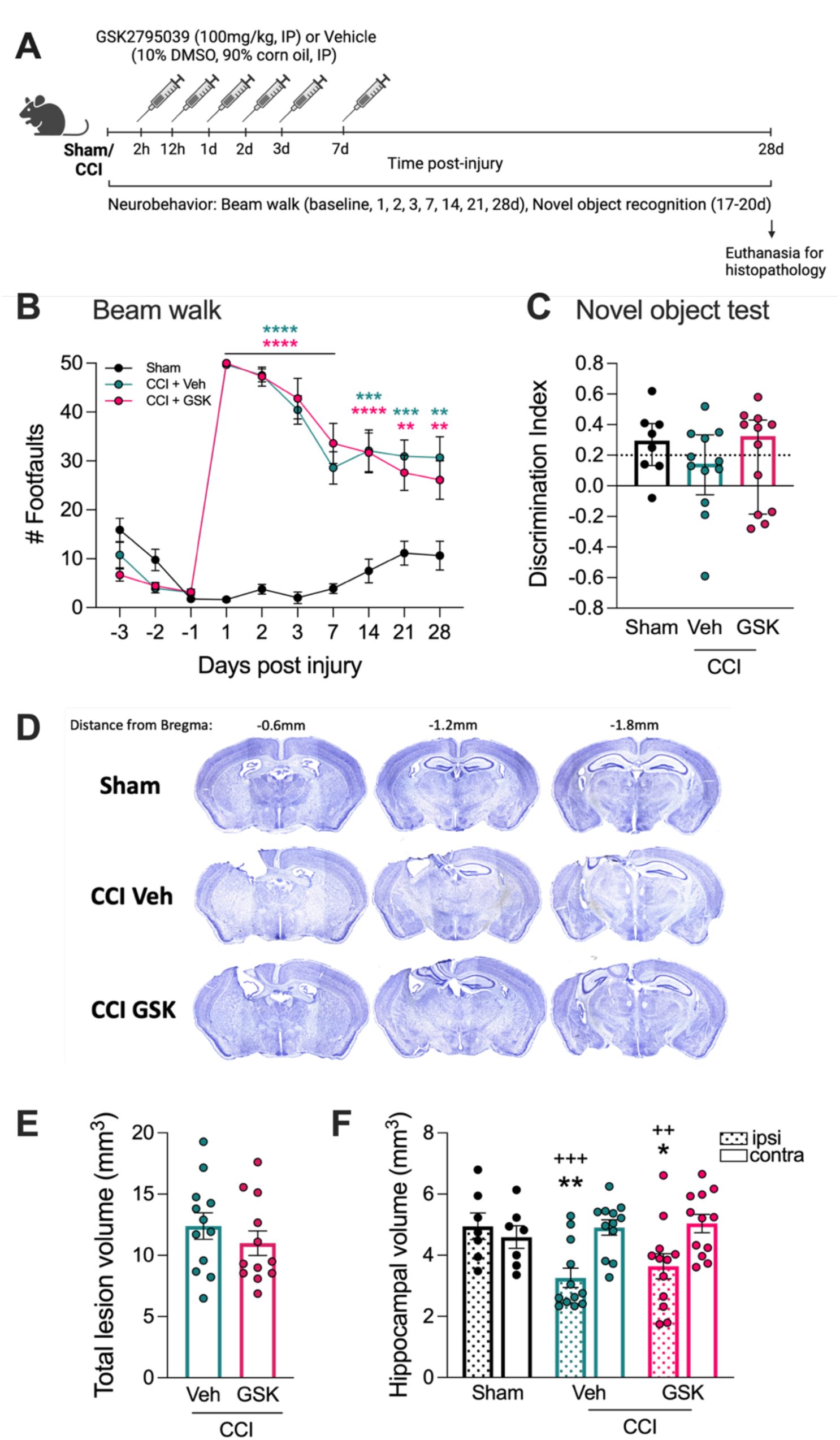
Pharmacological NOX2 inhibition fails to improve long-term neurological recovery following TBI but minorly reduces chronic neuropathology. GSK2795039 was administered at 2h, 12h, 1d, 2d, 3d and 7d post CCI (A). Fine motor coordination was assessed longitudinally by beam walk (A). Recognition memory was assessed by novel object recognition (NOR) test at 17-20 DPI (A). Cortical lesion and hippocampal volumes were calculated at 28 DPI. CCI, irrespective of treatment, induced increased number of foot-faults from 1-28 DPI (B; **p<0.01; ***p<0.001; ****p<0.0001 vs sham). GSK2795039 treated CCI mice showed a minor reduction in the number of foot-faults compared to vehicle treated, but levels failed to reach statistical significance (B). In the NOR test, CCI mice had reduced DI compared to sham, but levels failed to reach statistical significance (C). GSK2795039 treated CCI mice showed improved DI, but levels did not reach significance (C). CCI induced a total cortical lesion and GSK2795039 treatment showed reduced lesion volume, but levels failed to reach statistical significance (D, E; p=0.1755 vs CCI veh). CCI reduced ipsi hippocampal volume (F; *p<0.05; **p<0.01 vs sham ipsi; ^++^p<0.01; ^+++^p<0.001 vs contra within group). GSK2795039 treated CCI mice were statistically less different from sham than vehicle treated. Data are mean ±SEM (n=8-12 per group). **p<0.01; ***p<0.001; ****p<0.0001 vs sham; *p<0.05; **p<0.01 vs sham ipsi; ^++^p<0.01; ^+++^p<0.001 ipsi vs contra within group; Two-Way ANOVA with repeated measures and uncorrected Fisher’s LSD (B); Kruskal-Wallis test with Dunn’s multiple comparisons test vs CCI veh (C); unpaired One-tailed t-test (E); Two-Way ANOVA with Šidák’s multiple comparisons test (F).

When evaluating gross neuropathology at 28 DPI, vehicle treated CCI mice had a large cortical lesion that expanded into the ipsilateral hippocampus (**Figure 10D,E**; Veh + CCI = 12.4 ± 1.0 mm^3^). GSK2795039 treated CCI mice had a reduced lesion (GSK2795039 + CCI = 10.9 ± 1.0 mm^3^), although this failed to reach statistical significance. We finally checked if NOX2 inhibition could preserve hippocampal volume following injury. Vehicle treated CCI mice had reduced hippocampal volume in the ipsilateral (ispi) hemisphere compared to sham, and the contralateral (contra) side within group (**Figure 10D,F**; p<0.05; p<0.01 vs sham ipsi; p<0.01; p<0.001 vs contra within group). Although no statistically significant difference between Vehicle and GSK2795039 treated CCI groups was observed, GSK2795039 treated CCI mice were statistically closer to sham when compared to vehicle treated CCI mice.

## Discussion

NOX2, the phagocyte NADPH oxidase, contributes to age-related neurodegenerative diseases [1], and it has been demonstrated to drive persistent microglial activation and associated neurodegeneration up to 1 year post-TBI [2, 4, 5, 49]. Although there is increased recognition of the central role NOX2 plays in post-traumatic oxidative stress and neuroinflammation, pharmacological inhibition of NOX2 has been less well studied [50, 51]. Here, we investigated how a small molecule brain-penetrant NOX inhibitor (GSK2795039) with high selectivity for NOX2 over other isoforms (e.g. NOX4 etc.) [25] modulates the NOX2/ROS/NLRP3 inflammasome axis in microglia and in an experimental model of TBI in mice.

Using *in vitro* models of microglial-mediated neuroinflammation, we showed that NOX2 inhibition by GSK2795039 dose-dependently attenuated ROS signalling and NLRP3 inflammasome induced pro-inflammatory cytokine production (IL-1β, IL-18), pyroptotic cell death, as well as nitrite and TNF⍺ production, which are key mediators of microglial inflammatory response [40]. Next, we found that moderate-severe CCI in male mice increased NOX2/ROS/IL-1β+ expression in brain resident microglia and also dramatically increased the infiltration of inflammatory neutrophils and monocytes with the NOX2/ROS/IL-1β+ signature into the injured brain parenchyma. There was a dynamic temporal response, such that immune populations at different timepoints played a crucial role in promoting NOX2 and NLRP3 inflammasome activation after TBI. Microglial activation peaked at 3 DPI and persisted through 7 DPI with increased NOX2/ROS and IL-1β production. In contrast, infiltrating CD11b+CD45^hi^ myeloid cells were short-lived and there was few infiltrating NOX2/ROS/IL-1β+ cells in the injured brain at 7 DPI. This is consistent with literature that showed even though the number of infiltrating cells is reduced by 7 DPI, these myeloid cells have high ROS expression [34], which may be contributing to the evolving chronic neuroinflammatory response following TBI. Notably, here we demonstrated that systemic administration of the small molecule NOX2 inhibitor, GSK2795039, starting at 2h post-injury attenuated NOX2/ROS and NLRP3 inflammasome mediated activation of microglia, and to an even greater extent in infiltrating myeloid cells.

Not only does phagocyte NOX2 contribute to increased neuroinflammation it also initiates NLRP3 inflammasome activation by providing a second signal (i.e. ROS) for NLRP3 inflammasome assembly and IL-1β/IL-18 processing [11]. In a model of penetrating ballistic-like brain injury in rats there was increased NLRP3 inflammasome protein expression (e.g. ASC), IL-1β/IL-18, and GSDMD, indicative of the formation of the pyroptosome at 3 DPI [52]. Moreover, upregulated microglial ASC and IL-1β was observed up to 12 weeks post-injury [52]. In another study, delayed depletion of microglia using the CSF1R inhibitor, PLX5622, alleviated both NOX2 and NLRP3 inflammasome-associated neuroinflammation following TBI with reduced microglial caspase-1 activity and IL-1β release at 12 weeks post-injury [9]. Changes in microglial NOX2/NLRP3 inflammasome activation patterns were associated with improved motor and cognitive function recovery and reduced grey and white matter degeneration [9]. Notably, NOX2 knockout mice have reduced NLRP3 inflammasome activation acutely after TBI as shown by reduced ASC and cleaved IL-1β protein in the injured cortex, which was also associated with decreased lesion volume and neuronal loss [15].

The potent NLRP3 inhibitor, MCC950, has shown promise in preclinical studies to reduce TBI-induced neuroinflammation and associated neurodegeneration [45]. Post-injury systemic administration of MCC950 to TBI mice resulted in reduced NLRP3 inflammasome activation and microglial IL-1β expression in the injured cortex, preserved blood brain barrier (BBB) function, reduced lesion volume, and improved neurological deficits, including improved motor and cognitive function recovery after TBI [27]. Interestingly, the neuroprotective effect of MCC950 were completely abolished in injured mice that were depleted of microglia/macrophages using CSF1R inhibitor, PLX3397 [27]. Inhibitory effects on ROS-NLRP3 inflammasome interactions have been implicated in the neuroprotective actions of MCC950. The oxidative stress regulator, thioredoxin interacting protein (TXNIP), can directly associate with NLRP3 to induce inflammasome oligomerization and activation leading to cytokine production and proapoptotic signaling [53, 54]. TXNIP is upregulated in response to TBI [26] coincident with NLRP3 inflammasome activation, while MCC950 treatment robustly reduced TXNIP expression after TBI [26]. Mechanistically, this likely occurs downstream of ROS inhibition, which, in turn, promotes TXNIP expression through transcription factor forkhead box O3 (FoxO) in aged brains [55]. Our *in vitro* studies reveal the importance of the NOX2/ROS/NLRP3 inflammasome axis in microglial pro-inflammatory activation. Both the small molecule NOX2 inhibitor, GSK2795039, and the NLRP3 inhibitor, MCC950, robustly suppressed IL-1β/IL-18 production and microglial pyroptosis, demonstrating the inhibitory potential of suppressing NOX2 upstream of the NLRP3 inflammasome to reduce microglial-mediated neuroinflammation.

An important finding in our study was that GSK2795039 treatment had an inhibitory effect on NOX2/IL-1β+ infiltrating myeloid cells following TBI, which was more robust than in the microglial population. We, and others, have shown that blood borne CCR2+ inflammatory monocytes infiltrated the brain between 12 and 24 hours post-TBI and persisted for up to 7 days [4, 44, 56]. Furthermore, CCR2 antagonism blocks monocytes infiltration and alters the lesion microenvironment by reducing NOX2 expression [44, 56], which can reduce motor and cognitive deficits induced by TBI [44, 56]. Mechanistically, it has been shown that constitutive NOX2 knockout repolarizes peripheral macrophages toward an anti-inflammatory phenotype by increasing IL-10/STAT3 signalling, which reduces neuroinflammation and neurodegeneration in TBI mice [8]. Of note, GSK2795039 treatment had profound effects on NOX2/IL-1β expression in infiltrating neutrophils. Following TBI, ROS generated by NOX2 in neutrophils is known to contribute to oxidative stress and neuronal injury, and increased NOX2 activity facilitates release of matrix metalloproteinases (e.g. MMP-9), which degrade extracellular matrix to disrupt the BBB [57]. Upregulated NOX2/ROS in neutrophils can also activate other immune cells and enhance the expression of pro-inflammatory cytokines (e.g. IL-1β) [57]. Notably, when one of the cytosolic components of NOX2, neutrophil cytosolic factor 1 (NCF1), is constitutively knocked out in mice neutrophil infiltration into the injured brain parenchyma is reduced and neutrophil ROS levels are greatly suppressed [58]. NCF1 deficiency also results in reduced proinflammatory cytokine release at 3 DPI, including IL-1β, and protects against long-term motor coordination deficits following TBI [58]. Therefore, NOX2 inhibition in both infiltrating neutrophils and inflammatory monocytes can have profound effects on the evolving neuroinflammatory response in the TBI brain. As such, therapeutic strategies that interfere with NOX2/ROS in both brain resident (i.e. microglia) and infiltrating (i.e. neutrophils and monocytes) mononuclear cells may be required to improve overall TBI outcomes.

NOX2 inhibition also reduced the absolute numbers of CD3+ T cells and IL-1R expression on CD4+ and CD8+ T cells, indicating that myeloid-T cell crosstalk may be altered by post-injury GSK2795039 treatment. Interestingly, inhibition of NLRP3 inflammasome by MCC950 treatment also reduced the numbers of infiltrating CD4+ and CD8+ T cells acutely after TBI [27]. NOX2/NLRP3 inflammasome inhibition may alter myeloid cells-T cell crosstalk because of reduced IL-1β-producing myeloid cells and a simultaneous reduction in the number of IL-1R-expressing T cells. Indeed, brain resident and infiltrating innate immune cells recruit T cells during acute brain injury. It has been demonstrated that microglia drive transient insult-induced brain injury by chemotactic recruitment of CD8^+^ T lymphocytes in radiation-induced brain injury (RIBI) patients and mouse models [59]. Further, granzyme B-producing CD8+ cytotoxic T cells have been shown to drive long-term neurological impairment after TBI in mice [60], while CD8+ T cell-deficient mice promote an anti-inflammatory, IL-13, response. Not only does neutralization of CD8+ T cell activation alleviate neuroinflammation, but it also augments T regulatory (Treg) cell pro-reparative responses. IL-2 recruits Treg cells to the injured brain and regulates the neuroimmune response [61]. Notably, gene-delivery of IL-2 to the injured brain resulted in a reduced lesion size and improved cognitive function performance in IL-2 treated TBI mice [61]. These findings highlight the complexity of the neuroimmune milieu in injured tissue.

One of the ways GSK2795039 may be influencing infiltrating T cell populations *in vivo* is that NOX2 inhibition interferes with antigen presentation and thereby alters T cell recruitment, which has downstream effects on T cell-myeloid interactions. It has been shown in experimental autoimmune encephalomyelitis (EAE), that NOX2 expressing dendritic cells regulate and support myelin oligodendrocyte protein (i.e. MOG) antigen processing and presentation to T cells [62]. Knockout of *Cybb,* the gene encoding NOX2, specifically in dendritic cells (using *Itgax^Cre/+^Cybb*^fl/fl^ mice) restricted T cell recruitment into the CNS and ameliorated EAE disease development as demonstrated by reduced immune invasion, demyelination, and axonal damage [62]. Moreover, in post-mortem Multiple Sclerosis tissue there is increased NOX components, Cyba (p22^phox^) and Cybb (NOX2), in CNS resident (predominantly microglia) and infiltrating cells, particularly at the site of tissue damage and at perivascular areas near T cells [63]. Combined, these studies shed a light on the pathogenic role of the adaptive immune system in the injured CNS and crosstalk with microglia and infiltrating myeloid cells.

This preclinical study has several limitations, particularly related to GSK2795039 treatment. Hirano et al., demonstrated that NOX2 inhibition was lost within 24h following systemic administration of GSK2795039 as shown by undetectable GSK2795039 levels in the blood as well comparable ROS levels following paw inflammation in both GSK2795039 and vehicle treated mice at 24h [25]. Our pharmacokinetic analysis demonstrated therapeutically relevant GSK2795039 drug levels in the injured brain. A high drug clearance rate was observed with the concentration of GSK2795039 almost 5-fold greater in the liver compared to the brain, perhaps due to GSK2795039 being metabolised by liver microsomal fractions and preventing the dehydrogenation of the methylindoline moiety to reduce rodent metabolism [64]. As a result, we chose to administer GSK2795039 every 12-24h starting at 2h post-injury, until the day of sacrifice (in the 3DPI study) or up to 7 DPI (in the long-term 28 DPI study), to model a clinically relevant TBI scenario. The high drug clearance rate along with its short half-life in mice [25], may be why previous studies required repeated daily dosing of GSK2795039 to see cognitive improvements at 14 DPI in a rat TBI model [65]. Furthermore, others administered GSK2795039 prophylactically to ensure drug was in the system prior to TBI for proof-of-concept biochemical studies [66]. In a mouse model of spared-nerve injury, GSK2795039 was administered twice daily for 2 days starting at 1h prior to surgery, which resulted in reduced microglial activation at 2 DPI, but not at 11 DPI [67]. In our pre-clinical study, anti-inflammatory and neuroprotective responses following TBI were observed in the acute phase (at 3 DPI) but were lost in the chronic phase (at 28 DPI), indicating the effect of NOX2 inhibition was not sustained perhaps due to high drug clearance rates and suboptimal brain penetration. The drug dosing regimen of GSK2795039 for TBI and alternative routes of administration should be investigated in future studies to guarantee optimal drug efficacy. Another significant limitation of our study was that only male mice were used. Sex differences in neuroimmune response during the acute phase post-injury have been documented [34, 68] that include changes in NOX2/ROS and IL-1β levels in brain resident and infiltrating immune cells. Thus, it will be important to investigate the effects of GSK2795039 treatment in female mice to determine if similar anti-inflammatory effects are observed and/or the drug’s effect size.

To conclude, despite its importance, the highly complex neuroimmunological response to TBI remains under-investigated in the field. Addressing the many gaps in understanding of the role of innate and adaptive immunity in TBI is essential for future therapy development. Our findings identify the NOX2-ROS-NLRP3 inflammasome axis along with myeloid-T cell crosstalk as effective targets in the neuroinflammatory response following TBI. Our translational studies indicate that the small molecule NOX2 inhibitor, GSK2795039, may be a promising therapeutic drug for mitigating post-traumatic neuroinflammation and associated neurodegeneration, both in brain resident microglia, and peripheral immune cells that traffic to the injured brain, but further preclinical studies are needed.

## Acknowledgments

The authors thank Professor Kingston Mills and his group, particularly Dr. Charlotte Leane and Dr. Barry Moran for guidance and advice with flow cytometry. We thank Katherine Falcon for her help with hippocampal volume analysis.

---

**Supplemental Figure 1:**
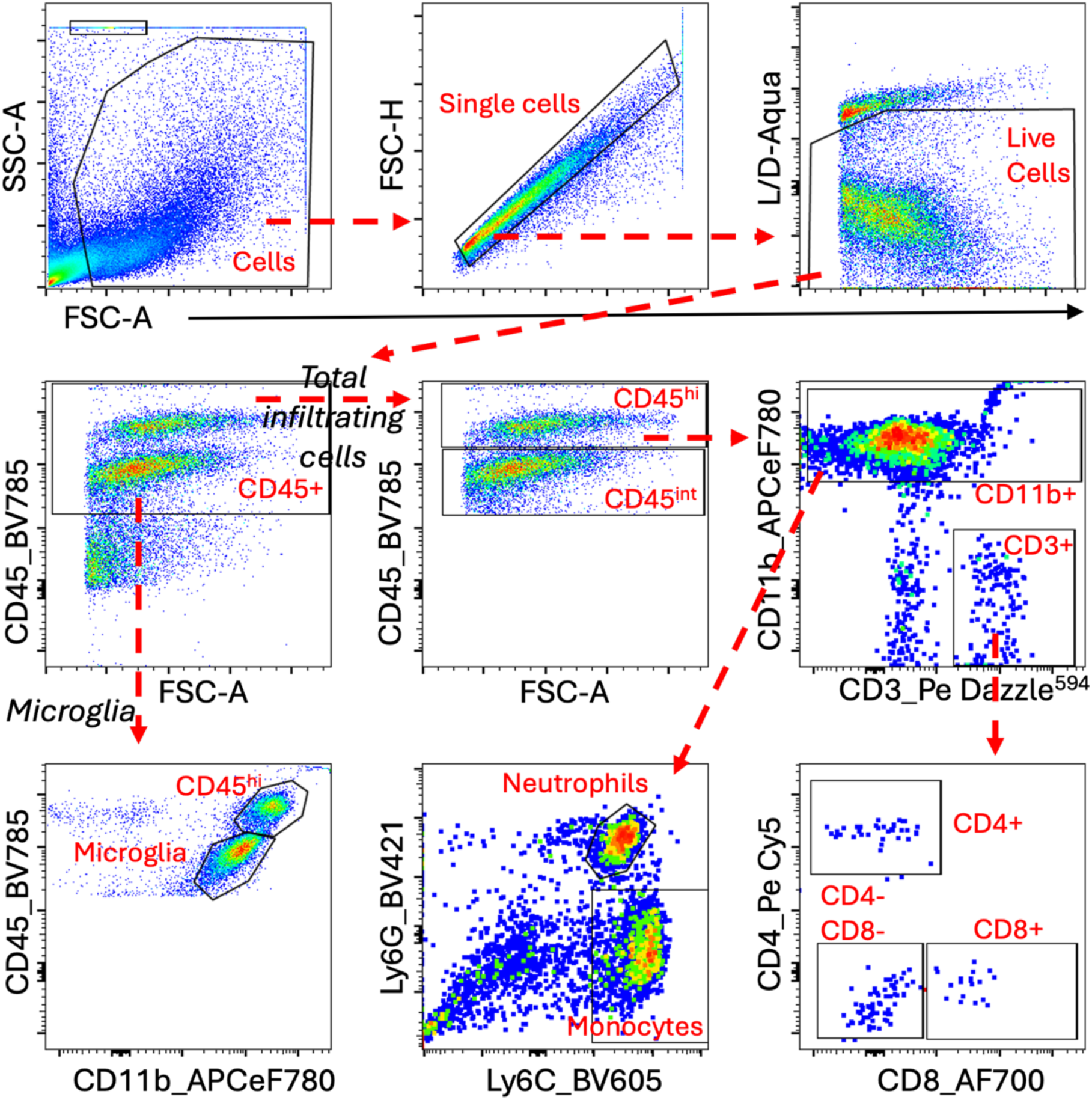
Flow cytometry gating strategy for brain. Microglia, neutrophils, monocytes and T cells in the brain were identified using the following gating strategy: CD45^+^ live cells consists of resident microglia (CD11b^+^CD45^int^) and infiltrating peripheral cells (CD11b^+^CD45^hi^), which include neutrophils (Ly6G^+^), monocytes (Ly6C^+^) and CD11b^-^CD3^+^ T cells (CD4^+^ and CD8^+^) in brain at 3 DPI. Abbreviations: FSC-A, forward scatter area; SSC-A, side scatter area; hi, high; int, intermediate; L/D, live/dead.

